# The efficacy of chemotherapy is limited by intratumoural senescent cells that persist through the upregulation of PD-L2

**DOI:** 10.1101/2022.11.04.501681

**Authors:** Selim Chaib, José Alberto López-Domínguez, Marta Lalinde, Neus Prats, Ines Marin, Kathleen Meyer, María Isabel Muñoz, Mònica Aguilera, Lidia Mateo, Camille Stephan-Otto Attolini, Susana Llanos, Sandra Pérez-Ramos, Marta Escorihuela, Fatima Al-Shahrour, Timothy P. Cash, Tamara Tchkonia, James L. Kirkland, Joaquín Arribas, Manuel Serrano

## Abstract

Anti-cancer therapies often result in a subset of surviving cancer cells that undergo therapy-induced senescence (TIS). Senescent cancer cells strongly modify the intratumoural microenvironment favoring immunosuppression and, thereby, tumour growth. An emerging strategy to optimise current therapies is to combine them with treatments that eliminate senescent cells. To this end, we undertook an unbiased proteomics approach to identify surface markers contributing to senescent cells immune evasion. Through this approach, we discovered that the immune checkpoint inhibitor PD-L2, but not PD-L1, is upregulated across multiple senescent human and murine cells. Importantly, blockade of PD-L2 strongly synergises with genotoxic chemotherapy, causing remission of solid tumours in mice. We show that PD-L2 inhibition prevents the persistence of chemotherapy-induced senescent cells, which exert cell-extrinsic immunomodulatory actions. In particular, upon chemotherapy, tumours deficient in PD-L2 fail to produce cytokines of the CXCL family, do not recruit myeloid-derived suppressor cells (MDSCs) and are eliminated in a CD8 T cell-dependent manner. We conclude that blockade of PD-L2 improves chemotherapy efficacy by reducing the intratumoural burden of senescent cells and their associated recruitment of immunosuppressive cells. These findings provide a novel strategy to exploit vulnerabilities arising in tumour cells as a result of therapy-induced damage and cellular senescence.

## Introduction

The inhibitory receptor PD-1 and its ligand PD-L1 (B7-H1) constitute an important immune checkpoint that controls the establishment of immune tolerance and negatively regulates the activity and proliferation of PD-1-expressing immune cells^1^. They are also key contributors to immune evasion by cancer cells, which frequently overexpress PD-L1^2^. Immunotherapies targeting PD-1/PD-L1 have been successfully used in the clinic against a broad spectrum of tumours^3-5^, including melanoma and non-small cell lung cancer^6^. An alternative PD-1 ligand, PD-L2 (B7-DC) has received comparatively less attention due to the lower frequency of PD-L2 positive cancers, compared to PD-L1^7^.

PD-L1 and PD-L2 share roughly a 40% of their amino acid sequence and they compete for PD-1 binding, although, interestingly, PD-L2 exhibits a 2- to 6-fold higher binding affinity to PD-1^8^. PD-L2 is predominantly expressed by dendritic cells, macrophages, and other antigen presenting cells, together with other immune populations. Additionally, in the context of cancer, cancer-associated fibroblast (CAF) can express PD-L2 and contribute to an immunosuppressive tumour microenvironment (TME)^9,10^. PD-L2 expression is most frequent in triple-negative breast cancer (TNBC) and abundant in small intestine and pancreatic neuroendocrine tumours, gallbladder cancer, endometrial cancer, bladder cancer, gastric cancer and head and neck squamous cell carcinoma (HNSCC)^7,11^. There is little understanding about the biological role of PD-L2 for immune evasion in cancers, although it is interesting to note that ectopic expression of PD-L2 produces immune evasion through inhibition of PD-1^12^. In the case of monocytes and macrophages, interferon γ (IFN-γ) upregulates both PD-L1 and PD-L2, while IL-4 selectively upregulates PD-L2^13,14^. However, at present, there is limited information regarding how levels of PD-L2 are regulated in cancer. Two recent reports have found upregulation of PD-L2 in some cancer cell lines in vitro in response to cisplatin^15^ or radiation^16^.

Conventional chemotherapy or radiotherapy is still the most common treatment for solid cancers. DNA damage and other insults associated with these therapies can trigger therapy-induced senescence (TIS) in cancer cells and in other intratumoural cells, such as endothelial cells and fibroblasts^17^. The senescence program constitutes a cell-intrinsic barrier against oncogene-driven proliferation and transformation, however, it also involves a prominent secretory activity known as the Senescence Associated Secretory Phenotype (SASP)^18^. The SASP is complex and heterogeneous, comprising pro-inflammatory cytokines, chemokines, matrix remodeling enzymes and growth factors, that result in cell-extrinsic effects known to facilitate tumour growth^19-23^. More specifically, the SASP includes immunosuppressive factors, like TGF-β, and chemokines that recruit myeloid-derived suppressive cells (MDSCs)^24-27^. Therefore, while senescence is a cell-intrinsic barrier for cancer cells, it is also pro-tumourigenic through its extrinsic actions on the tumour microenviroment.

Therefore, therapies that selectively eliminate senescent cells (known as senolytic therapies) have been demonstrated in multiple studies to synergise with cancer chemotherapy^28^. This is the case of senolytic compounds, such as ABT-263 (navitoclax)^23^, derivatives of this drug^29,30^, dasatinib^31,32^, cardiac glycosides^33,34^ and mTOR inhibitors^35^. Also, elimination of senescent cells using engineered CAR-T cells leads to improved tumour control in combination with MEK and CDK4/6 inhibitors^36^. Whether PD-L2 plays a role in the senescent phenotype and whether this feature can be exploited in novel therapies is unknown. Here, we show the importance of PD-L2 upregulation in the persistence of therapy-induced senescent cells and, thereby, in their ability to modulate tumour immunosurveillance across a variety of mouse models of cancer.

## Results

### Senescent cancer cells overexpress PD-L2

To gain insight into the immunomodulatory potential of senescent cells, we performed an unbiased proteomic screen of proteins enriched in the plasma membrane of senescent and non-senescent human SK-MEL-103 melanoma cells, using doxorubicin or the CDK-4/6 inhibitor palbociclib to trigger therapy-induced senescence (TIS). Interestingly, among the upregulated plasma membrane proteins in both TIS conditions (doxorubicin and palbociclib) was PD-L2 (**Supplementary Table 1**). Moreover, drug class enrichment analysis of datasets from the LINCS L1000 project^37^ revealed that PD-L2 is upregulated by common inducers of cellular senescence, such as DNA-damage and cell cycle inhibition (**Extended Data Fig. 1a**). Indeed, PD-L2 transcript levels were significantly upregulated in TIS in human and murine cancer cell lines of various origins, including melanoma, lung squamous cell carcinoma, head and neck squamous cell carcinoma, and osteosarcoma (**Fig. 1a-b** and **Extended Data Fig. 1b-d**). In contrast, increases in PD-L1 transcript levels were modest or absent. PD-L2 expression was also elevated in vivo following the induction of TIS in human xenografts and mouse syngeneic tumours (**Fig. 1c-d** and **Extended Data Fig. 1e**).

**Fig. 1.**
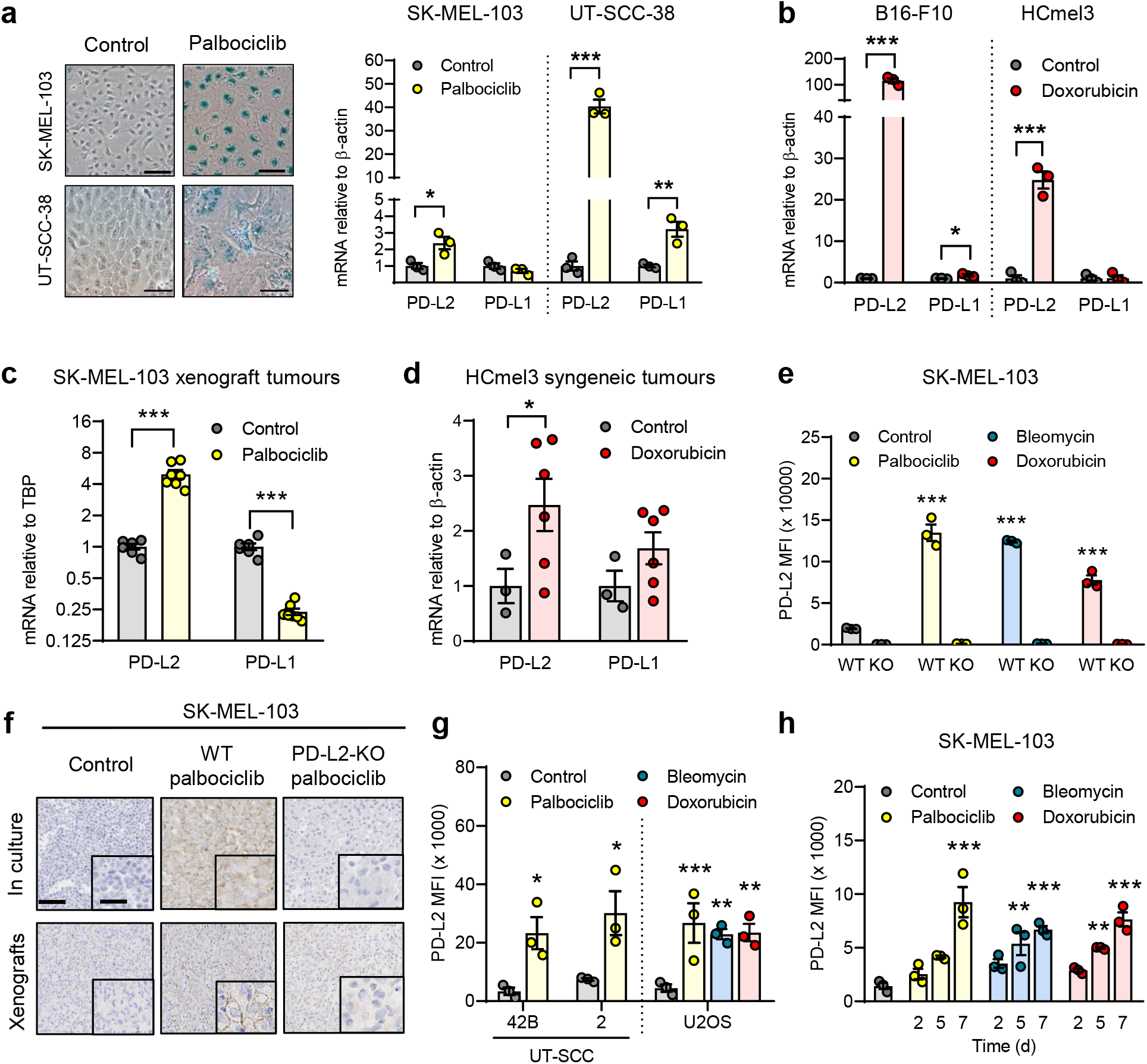
PD-L2 is upregulated in human and murine cancer cell lines upon induction of cellular senescence. (a) Representative images of senescence-associated b-galactosidase (SABG) staining and PD-L1/2 mRNA expression in human cancer cell lines treated with palbociclib. Scale bars = 50 µm. (b) PD-L1/2 mRNA expression in murine cancer cell lines after treatment with doxorubicin. (c) PD-L1/2 mRNA expression in SK-MEL-103 xenograft tumours in nude mice, treated with palbociclib. (d) PD-L1/2 mRNA expression in HCmel3 tumours in C57BL/6 mice treated with doxorubicin. (e) Quantification of PD-L2 protein levels by FACS upon generation of a PD-L2-KO SK-MEL-103 cell line, in control and senescent conditions. (f) PD-L2 stainings of WT and PD-L2-KO SK-MEL-103 cells in culture and as xenograft tumours, untreated and treated with palbociclib. Scale bars are 100 µm for low magnification and 50 µm for high magnification images. (g) PD-L2 protein levels as measured by flow cytometry upon induction of senescence with palbociclib, bleomycin and doxorubicin in different cancer cell lines at day 7 after induction. (h) PD-L2 protein levels as measured by flow cytometry upon induction of senescence with palbociclib, bleomycin and doxorubicin in SK-MEL-103 cells. MFI = median fluorescence intensity. t-tests or 1 way ANOVA with Tukey post-test were applied. * p < 0.05, ** p < 0.01, *** p < 0.001 vs respective control groups.

Based on the above findings, we next assessed cell surface levels of PD-L2 in TIS. Flow cytometry and immunohistochemistry-based analysis revealed elevated PD-L2 in SK-MEL-103 cells, both in vitro and in xenograft models, upon TIS (**Fig. 1e-f** and **Extended Data Fig. 1f**), which was not present in a PD-L2-KO SK-MEL-103 clonal cell line generated by CRISPR-Cas9 (**Extended Data Fig. 1g**). Upregulation of PD-L2 was detected by flow cytometry in cell lines of different cancer cell types that underwent TIS (**Figure 1g**) and occurred gradually during the development of TIS over a period of 7 days (**Fig. 1h**). It is interesting to note that osteosarcoma Saos-2 cells, which are unable to undergo senescence (due to the absence of RB1^38^), did not upregulate PD-L2 in response to palbociclib, in contrast to TIS-competent osteosarcoma U2OS cells (**Extended Data Fig. 1h**). Together, these data indicate that the upregulation of PD-L2 is a common feature of the program elicited by therapy-induced senescence (TIS) and results in elevated plasma membrane PD-L2 levels.

### PD-L2-KO tumours are highly susceptible to chemotherapy

The consistent upregulation of PD-L2 in therapy-induced senescent cells prompted us to study its relevance in the context of cancer therapy. For this, we generated a murine PD-L2-KO Panc02 cell line by CRISPR-Cas9 (**Extended Data Fig. 2a**) and we injected WT and PD-L2-KO cells orthotopically into the pancreas of immunocompetent mice that were subsequently treated with genotoxic chemotherapy (doxorubicin). Interestingly, chemotherapy controlled the growth of PD-L2-KO tumours significantly better than their WT counterparts (**Fig. 2a**). As a result, mice bearing PD-L2-KO tumours and treated with doxorubicin lived significantly longer than mice with treated WT tumours or untreated PD-L2-KO tumours (**Fig. 2b**). To extend our observations to a different tumour model, we injected WT and PD-L2-KO B16-OVA melanoma cells orthotopically into C57BL/6 mice. Again, we observed a profound reduction in tumour growth rate in PD-L2-KO tumours treated with chemotherapy (**Extended Data Fig. 2b**). In contrast, WT tumours had a partial and transient reduction in tumour growth in response to chemotherapy (**Extended Data Fig. 2b**). Collectively, these data indicate that the expression of PD-L2 by cancer cells limits the efficacy of chemotherapy.

**Fig. 2.**
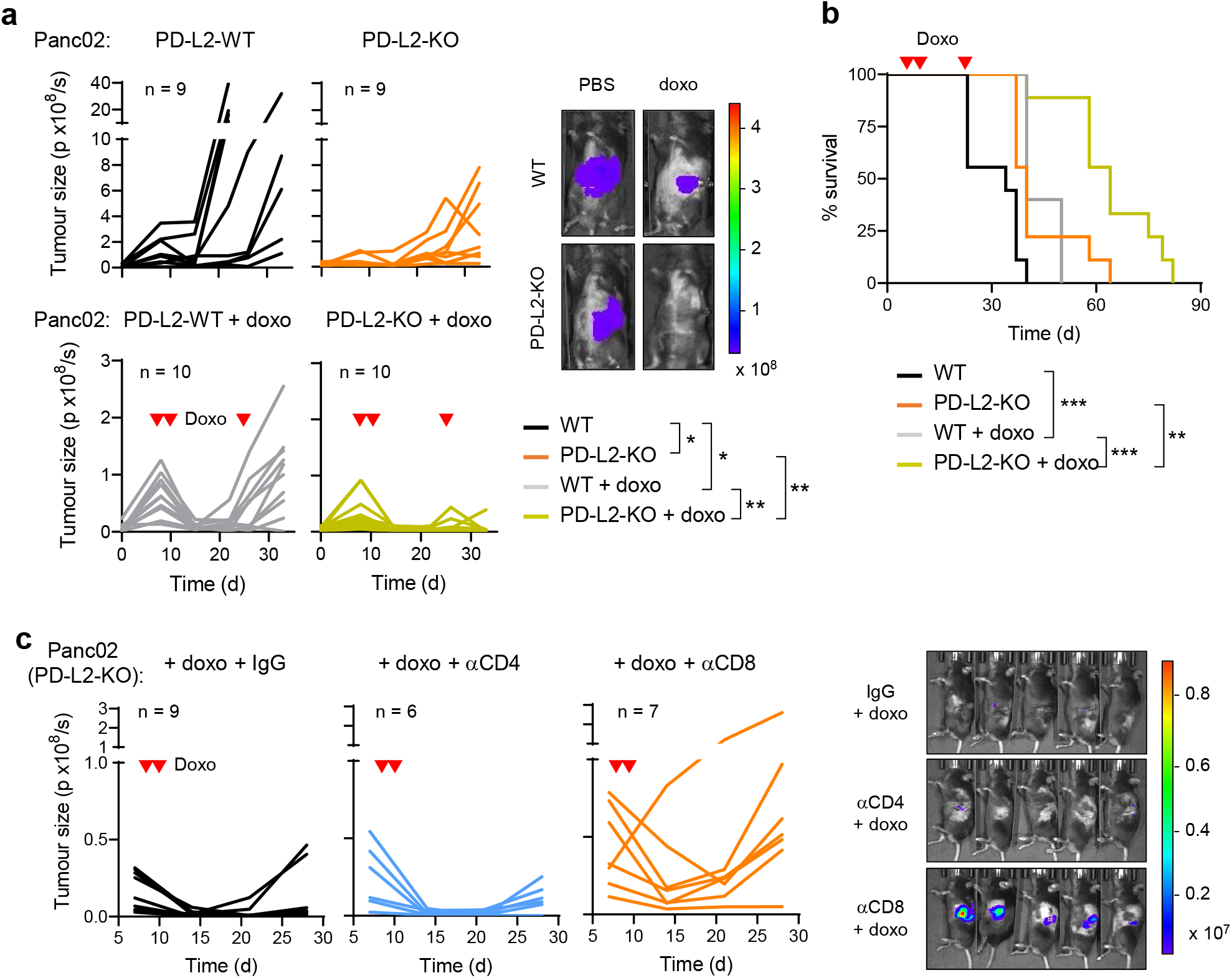
A combination of PD-L2 ablation and chemotherapy results in tumour remission. (a) Representative images and quantification of tumour growth for PD-L2-WT and KO Panc02 tumours, untreated or treated with doxorubicin at days 7, 10 and 24. (b) Survival curve for the mice from panel (a). (c) Quantification and representative images of tumour growth for PD-L2-KO Panc02 tumours after doxorubicin treatment (days 7 and 10) and repeated injections with IgG isotype control, anti-CD4 or anti-CD8 blocking antibody from day 3 after surgery. Luminescence units are photon/second (p/s) in the graphs and photon/sec/cm^2^/stereoradian in the images. 2-way ANOVA and 1 way ANOVA with Tukey post-test were used. * p < 0.05, ** p < 0.01, *** p < 0.001.

We next analysed whether a functional adaptive immune response was required for the control of PD-L2-KO tumours. For this, we injected WT and PD-L2-KO Panc02 cells into the pancreas of athymic nude mice (lacking T cells) and quantified tumour growth over time. Neither in the absence of chemotherapy nor after administration of doxorubicin (**Extended Data Fig. 2c**) did tumours show any significant difference in growth rate across experimental groups, suggesting that the adaptive immune system was responsible for the phenotype associated to PD-L2-KO tumours. To elucidate which major T cell subset was essential for tumour regression, we depleted CD4^+^ or CD8^+^ T cells from animals bearing PD-L2-KO pancreatic tumours and were subsequently treated with doxorubicin (**Extended Data Fig. 2d**). Mice lacking CD8^+^ T cells were unable to control PD-L2-KO tumours after chemotherapy, while control animals or mice treated with anti-CD4 blocking antibodies presented a robust suppression of tumour growth (**Fig. 2c**). Together, these results indicate that CD8^+^ T cells are responsible for the efficient removal of PD-L2-KO tumours upon chemotherapy.

### Absence of PD-L2 expression prevents the recruitment of myeloid cells after chemotherapy

To better understand the mechanisms underlying the enhanced efficiency of chemotherapy in PD-L2-KO tumours, we performed a comprehensive analysis of the immune infiltrate in Panc02 PD-L2 WT and KO tumours. We quantified a panel of sixteen markers of different immune populations by mass cytometry 5 days after the start of doxorubicin treatment (**Fig. 3a**). We detected a substantial recruitment of CD11b^+^Gr1^+^ myeloid cells in PD-L2-WT tumours after treatment with doxorubicin (**Fig. 3b**). This was in sharp contrast to PD-L2-KO tumours, which were unable to recruit these myeloid cells (**Fig. 3b**). No significant variations were observed across experimental conditions for other immune populations, including T cells, macrophages, or NK cells (**Extended Data Fig. 3a-c**). Interestingly, we also observed that depletion of CD8^+^ T cells resulted in accumulation of Gr1^+^ cells in chemotherapy-treated PD-L2-KO tumours (**Fig. 3c** and **Extended Data Fig. 3d**). To evaluate the impact of Gr1^+^ myeloid cells in this tumour model, we depleted Gr1^+^ cells from animals bearing WT tumours and treated with doxorubicin, observing an improvement in tumour control (**Fig. 3d**). These results indicate that, upon chemotherapy, immunosuppressive myeloid cells are recruited in a PD-L2-dependent manner. Conversely, absence of PD-L2 prevents recruitment of Gr1^+^ immunosuppressive myeloid cells post-chemotherapy, rendering tumours susceptible to the action of CD8^+^ T cells.

**Fig. 3.**
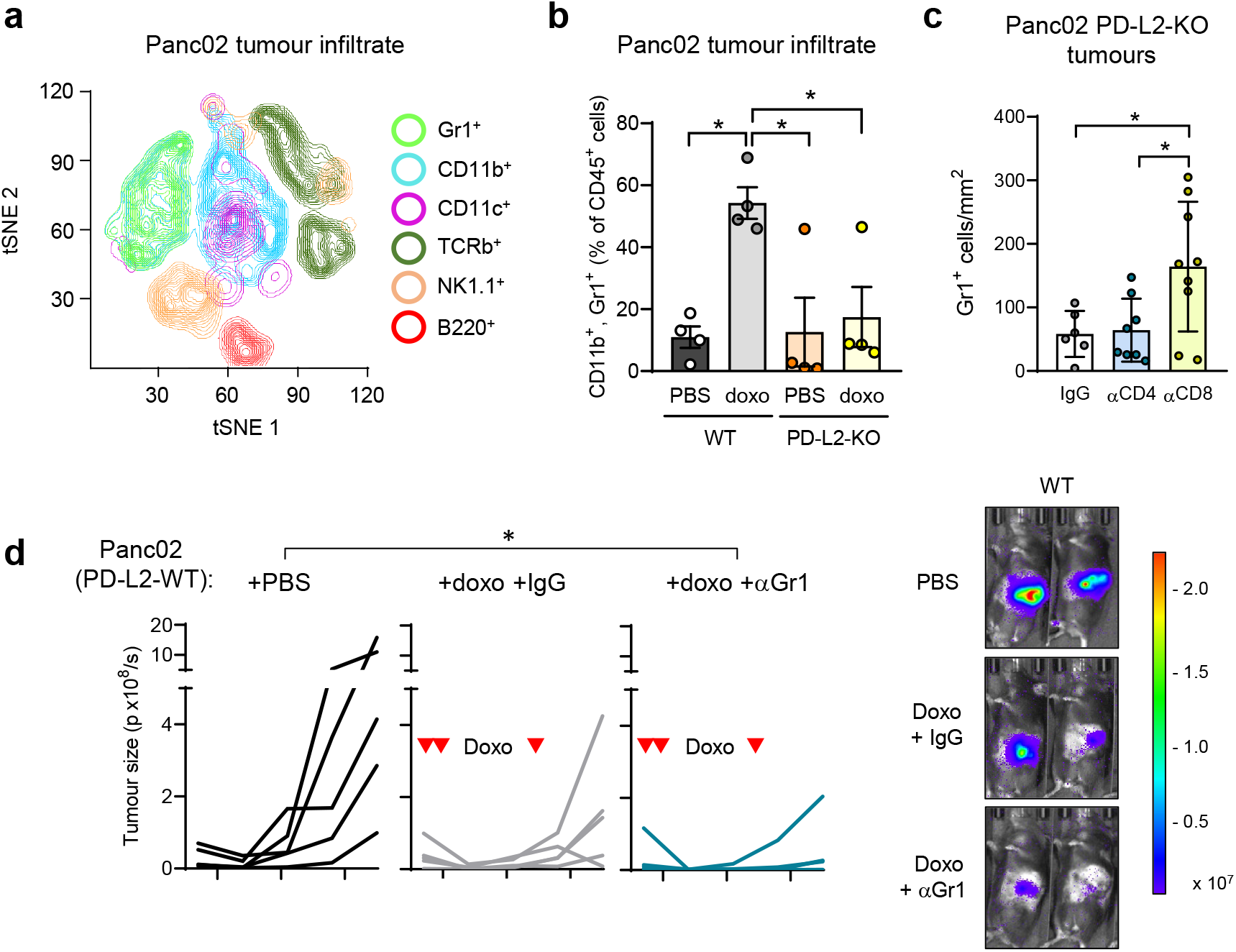
Recruitment of tumour-promoting myeloid cells is prevented in PD-L2-KO upon doxorubicin treatment. (a) t-SNE plot including the different tumour-infiltrating immune subpopulations detected by CyTOF. A pool of all 16 individuals in different experimental conditions is shown (n = 4 for WT and PD-L2-KO, untreated or treated with doxorubicin). Doxorubicin treatment was administered on days 7 and 10 and samples were obtained on day 12. (b) Quantification of the percentage of CD11b^+^ Gr1^+^ cells, relative to total CD45^+^ cells. N = 4 for each experimental group. (c) Quantification of Gr1^+^ cells in sections of PD-L2-KO tumours treated with doxorubicin on days 7 and 10, subject to depletion of CD4^+^ or CD8^+^ T cells and collected on day 28. (d) Representative images and quantification of tumour growth for PD-L2-WT tumours, untreated or treated with doxorubicin on days 7, 10 and 24, including an additional group treated with anti-Gr1 blocking antibody. N = 5 per group. Luminescence units are photon/second (p/s) in the graphs and photon/sec/cm^2^/steradian in the images. 1 way ANOVA with Tukey post-test was applied. * p < 0.05, ** p < 0.01, *** p < 0.001.

### PD-L2 determines the persistence of tumour senescent cells post-chemotherapy

We wondered about the degree of similarity between senescent cancer cells lacking or expressing PD-L2. To evaluate this, we analysed by RNAseq the transcriptome of both cell lines upon doxorubicin. Remarkably, their expression profiles were essentially identical with only 10 differentially expressed genes (DEGs) out of ∼21,000 detected genes (|FC| > 1.5, FDR < 0.05) (**Supplementary Table 2**). The enrichment of signatures of p53 activation and inflammation were evident and similar in WT and KO doxorubicin-treated Panc02 cells (**Extended Data Fig. 4**). These results suggest that PD-L2 is not relevant for the induction of senescence in vitro.

Given the high similarity between the transcriptomes of PD-L2 WT and KO cells, we wondered whether PD-L2 is important for the intratumoural persistence of senescent after chemotherapy. To evaluate senescent cell burden in orthotopic pancreatic tumours, we quantified senescence-associated beta-galactosidase (SABG) and p21^+^ cells by immunohistochemistry. Interestingly, 5 days post-chemotherapy, SABG^+^ and p21^+^ cells were clearly increased in PD-L2-WT tumours, but not in PD-L2 KO tumours (**Fig. 4a**). We interpret that, in accordance with our previous results, the absence of PD-L2 impairs the persistence of intratumoural senescent cells post-therapy.

**Fig. 4.**
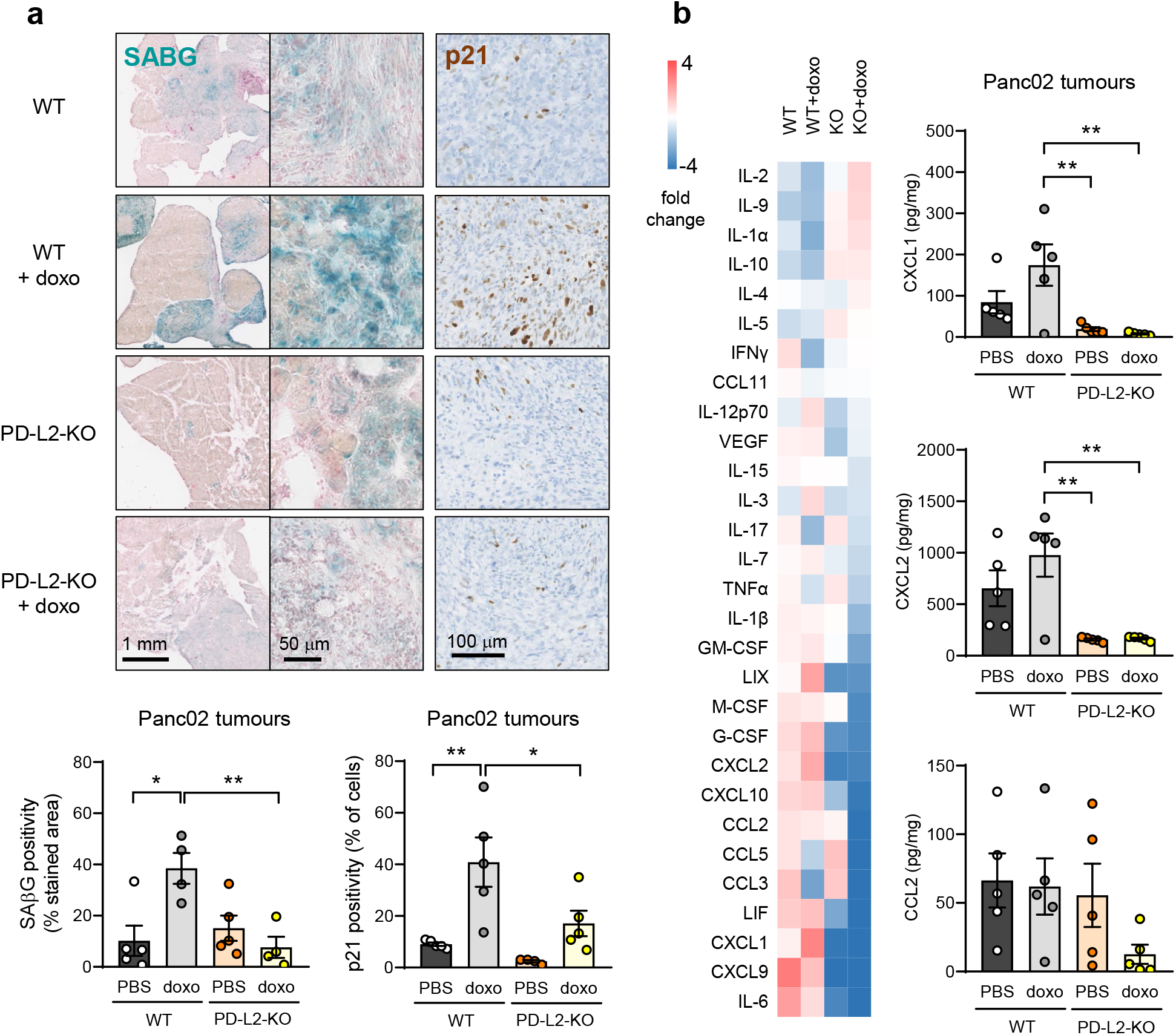
PD-L2-KO Panc02 tumours after chemotherapy present lower levels of cellular senescence and key cytokines and chemokines. (a) Representative images and quantification of SABG and p21 stainings in PD-L2-WT and KO, doxorubicin-treated (days 7 and 10) and untreated tumours, five days after the first dose. (b) Relative levels of intratumoural cytokines and chemokines measured by a commercial multiplexed system with beads bound to antibodies. Absolute quantifications of select cytokines and chemokines (CXCL1, CXCL2, CCL2). N = 5. 1 way ANOVAs with Tukey post-tests were applied. * p < 0.05, ** p < 0.01, *** p < 0.001.

It is known that senescent cells contribute to the generation of an immunosuppressive microenvironment by recruiting MDSC into the tumour^24-27^. A number of cytokines and chemokines present in the SASP mediate this recruitment, including IL-6^24^, CCL2^25^, CXCL1 and CXCL2^26^. We quantified the levels of intratumoural chemokines and cytokines in our Panc02 model and observed that cytokines associated with MDSCs recruitment were lowest in PD-L2-KO tumours post-chemotherapy, particularly in the case CXCL1 and CXCL2 (**Fig. 4b**). Our results suggest that lower levels of cellular senescence and decreased myeloid cell recruitment by secreted factors contribute to the synergic effect of chemotherapy and PD-L2 deficiency.

### Combinational therapy with blocking anti-PD-L2 antibodies

Given our previous results, we explored the role PD-L2 activity in additional models. MMTV-PyMT mice develop spontaneous mammary gland tumours, reaching the status of adenoma and carcinoma stages at 9 and 13 weeks of age, respectively. At 9 weeks of age, mice were weekly treated with doxorubicin for 4 weeks and tumours were analysed. In agreement with our previous models, we observed abundant senescent cells (SABG^+^) in post-therapy tumours (**Extended Data Fig. 5a**). Notably, SABG^+^ cells were also PD-L2^+^ by immunohistochemistry, further confirming that PD-L2 is a marker of therapy-induced senescence (**Extended Data Fig. 5a**). To determine the role of PD-L2 in this setting, we used a commercially available blocking anti-PD-L2 antibody (TY-25). Treatment with TY-25 alone did not have a significant effect on tumour growth and doxorubicin had a modest effect in reducing tumour growth rate (**Fig. 5a**). Remarkably, the combination of both treatments, TY-25 and doxorubicin, resulted in complete tumour regression (**Fig. 5a**). To characterise the anti-tumour effect of combined chemotherapy and PD-L2 antibody, we analysed lymphocytic infiltration by immunohistochemistry and observed a highly significant increase of CD8^+^ T cells, but not CD4^+^ T cells (**Fig. 5b-c**). To test the involvement of CD8^+^ T cells in the remission elicited by anti-PD-L2 treatment, we performed another experiment including anti-CD8 and anti-CD4 blocking antibodies. While blocking CD4^+^ cells had little effect, blocking CD8^+^ cells led to tumour regrowth (**Extended Data Fig. 5b**). Together, these results indicate increased immune surveillance of senescent and non-senescent tumour cells upon PD-L2 suppression.

**Fig. 5.**
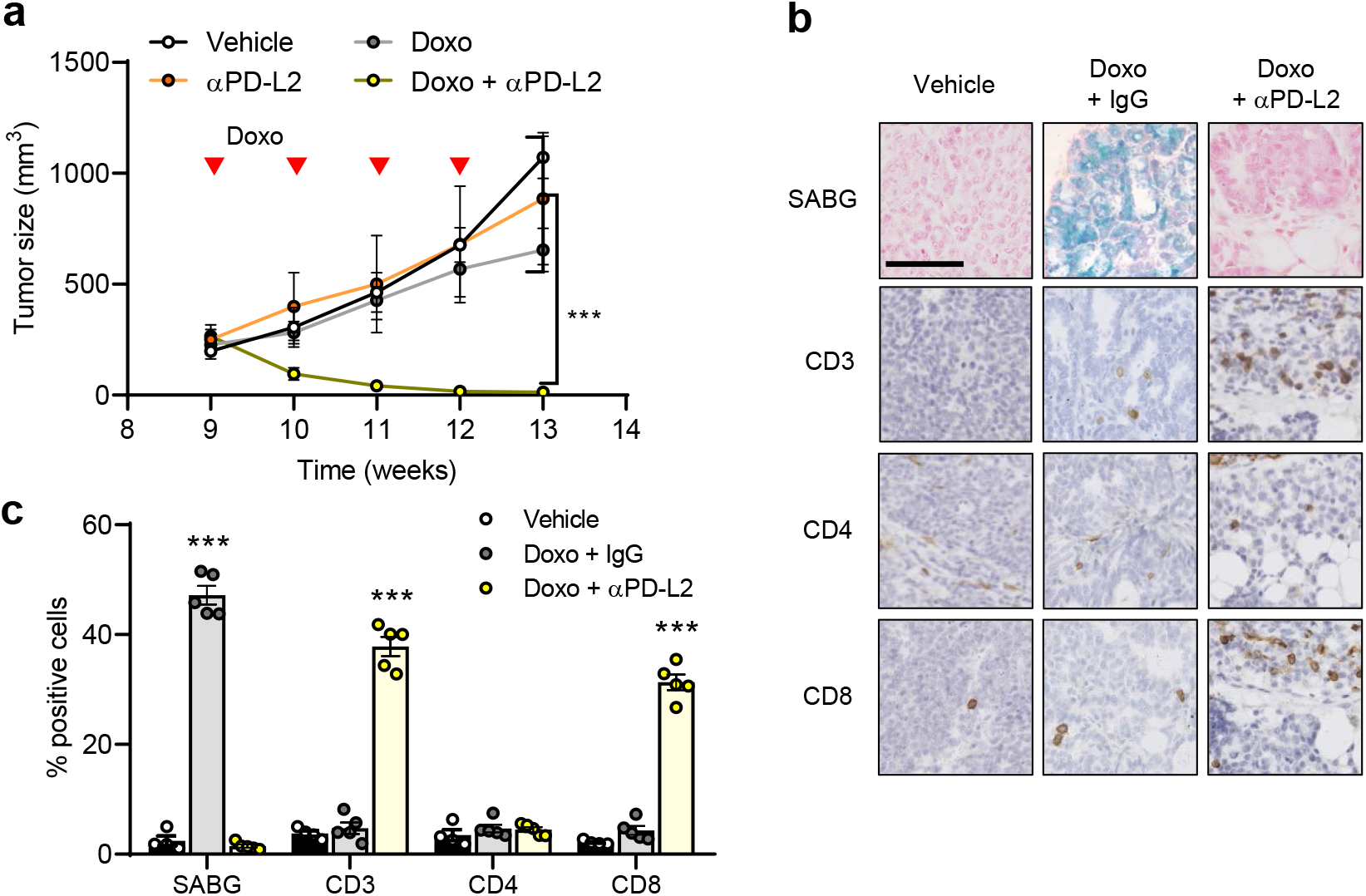
PD-L2 blockade successfully eliminates mammary tumours after chemotherapy. (a) Tumour growth in PyMT mice treated weekly (from week 9) with anti-PD-L2 alone (TY-25), doxorubicin alone, or a combination of both as indicated. 2-way ANOVA, *** p < 0.001. (b-c) SABG, CD3, CD4, and CD8 positivity in untreated tumours and tumours treated with doxorubicin alone or in combination with anti-PD-L2 blocking antibody. Representative images are shown. 1-way ANOVA with Tukey post-test. *** p < 0.001 vs the rest of experimental conditions. Scale bar = 100 µm.

## Discussion

Here, we demonstrate that senescence-inducing therapies employed in the clinic result in upregulation of PD-L2 in cancer cells. While the expression of PD-L2 is not necessary for senescence induction and maintenance, it contributes to immune evasion of post-therapy tumours. The presence of senescent cancer cells within tumours, even if they account to a subset of the tumor cells, is of high relevance due to their immunosuppressive secretome. In agreement with previous reports^24-27^, we show that PD-L2-proficient tumours secrete cytokines and chemokines, such as CXCL1 and CXCL2, which recruit myeloid-derived suppressive cells (MDSCs) upon therapy. Importantly, tumours deficient in PD-L2 fail to produce these chemokines and to recruit MDSCs, rendering tumours vulnerable to CD8^+^ T cells. We finally show that anti-PD-L2 antibodies unleash the immune clearance of tumours post-chemotherapy, including senescent as well as non-senescent cancer cells.

The accumulation of senescent cells with aging^39-41^ and in multiple pathological conditions^42-48^, including cancers^31^, has been partially attributed to defective immunosurveillance^49-55^. It remains to be explored to what extent PD-L2 is also involved in the accumulation of senescent cells associated to aging and aging-associated diseases.

The concept of immune-mediated senescent cell clearance being synergistic with senescence-inducing therapies has been recently reported in the context of cancer^36^. Specifically, in a model of pulmonary adenocarcinoma, a senescence-inducing therapy followed by administration of uPAR-specific CAR-T cells extended survival and increased lymphocytic infiltration^36^. Our current findings on the use of anti-PD-L2 antibodies add another strategy to improve the efficacy of anti-cancer therapies.

## Methods

### Mammalian tissue culture

SK-MEL-103 (human melanoma), U2OS (human osteosarcoma), Saos-2 (human osteosarcoma), HEK293T (human embryonic kidney) and H226 (human squamous cell carcinoma) cells were obtained from American Type Culture Collection. UT-SCC-38, UT-SCC-42B, UT-SCC-2 cells (human head and neck squamous cell carcinoma) were provided by Dr. Reidar Grenman (University of Turku, Finland). B16F1 and B16F10 cells (mouse melanoma) were provided by Dr. María Soengas (Spanish National Cancer Research Center, Spain). B16-OVA: B16 (mouse melanoma) expressing ovalbumin (OVA) and Panc02 (mouse pancreatic adenocarcinoma) were provided by Dr. Federico Pietrocola (Institute for Research in Biomedicine, Spain). HCmel3 cells (mouse melanoma) were provided by Dr. Thomas Tüting (University of Bonn, Germany). Cells were routinely tested for mycoplasma contamination. HcMel3 and Panc02 cells were maintained in Roswell Park Memorial Institute medium (Gibco). The rest of the cell lines were maintained in Dulbecco’s Modified Eagle’s Medium (Gibco). All media were supplemented with 10% fetal bovine serum (Gibco) with 1% penicillin/streptomycin (Gibco). All cell lines were cultured at 37° C in a humidified atmosphere and 5% CO_2_ and procedures were conducted under aseptic conditions in a biological safety cabinet according standard operating procedure.

### Induction of cellular senescence in tissue culture

Unless otherwise noted, senescence was induced using 5 μM palbociclib (Pfizer Inc.) for seven days, or 48 h treatments of 200 nM doxorubicin (Sigma-Aldrich) and 12 mU bleomycin (Sigma-Aldrich), after which fresh media was added. Senescence was evaluated at day 7.

### Gene expression analysis by qPCR

Total RNA from adherent cells or homogenised tissue biopsies was isolated using TRI Reagent (Sigma-Aldrich) according to manufacturer’s instructions. A total of 3-4 μg of total RNA was reverse transcribed using the iScript advanced cDNA synthesis kit (Bio-Rad). Quantitative PCR of target genes (Table 1) was performed using SybrGreen (Applied Biosystems) and ran on QuantStudioTM 6 Flex Real-Time PCR System using QuantStudioTM 6 and 7 Flex Real-Time PCR software v1.0 (Applied Biosystems). Relative gene expression levels were quantified using β-actin or human TBP as housekeeping genes, as indicated.

Primers used for human (h) and mouse (m) target genes

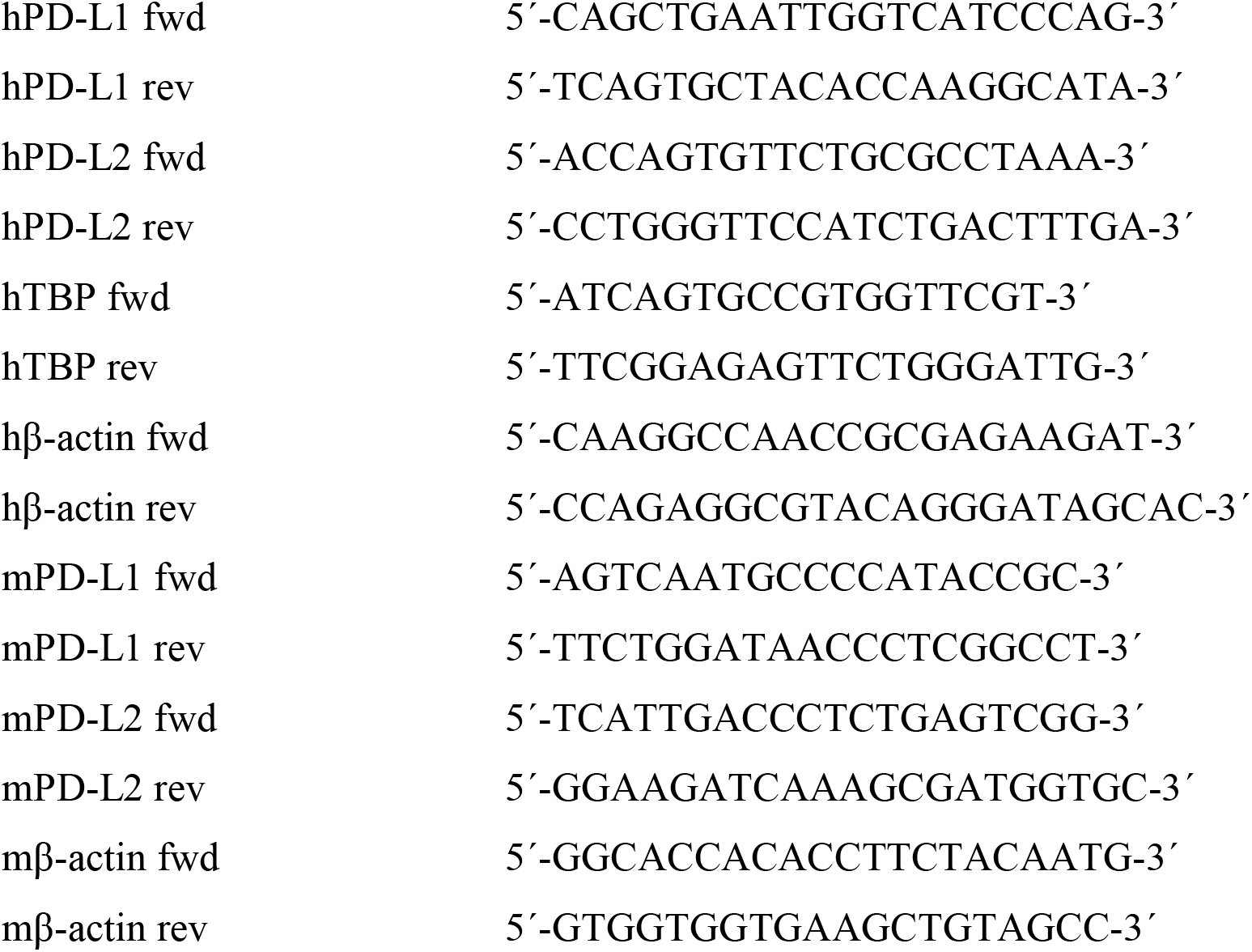

### Generation of cell pellets for immunohistochemistry

Cells were harvested using PBS + 10 mM EDTA at 37º C. Harvested cells were then centrifuged at 300 x g and washed with PBS. Cell pellets were then overlaid with 10% buffered formalin (Sigma-Aldrich) and fixed for 10 hours at 4º C. Formalin was then removed and cell pellets were embedded in paraffin. Tissue sections were deparaffinised, rehydrated and washed with EnVision FLEX wash buffer (Dako). Antigen retrieval was performed using Tris-EDTA buffer (pH 9) at 97º C for 20 minutes. After blocking endogenous peroxidase, slides were blocked with 5% goat serum + 2.5% bovine serum albumin (BSA) for 60 minutes. Slides were then incubated with anti-PD-L2 (CST #82723) diluted 1:25 in EnVision FLEX antibody diluent (Dako) over night at 4º C. Next day, slides were incubated with anti-goat-HRP for 45 minutes and developed for 10 minutes adding DAB. Slides were then dehydrated and mounted with DPX.

### Flow cytometry

Cells were harvested in PBS with 10 mM EDTA at 37º C. Collected cells were then stained with yellow live/dead dye solution (Invitrogen) for 15 minutes at 4º C and washed. Cells were then stained with anti-PD-L2-biotin (Miltenyi Biotec #130-098-525) diluted 1:11 in FACS buffer (0.5% BSA, 2 mM EDTA in PBS) for 15 minutes and after two washes the samples were incubated with anti-biotin-APCVio770 (Miltenyi Biotec #130-113-851), diluted 1:50 in FACS buffer for 15 minutes. After repeated washes, cells were filtered using a 70 μm cell strainer and analysed on a Gallios flow cytometer and with FlowJo 10.0.7.

### CRISPR

For PD-L2 knockouts, mouse and human sgRNAs were designed using the CHOPCHOP web tool: (http://chopchop.cbu.uib.no) and cloned into pSpCas9(BB)-2A-Puro (PX459) (Addgene #48139) and lentiCRISPRv2 (Addgene #52961).

sgRNAs targeting human (h) or mouse (m) PD-L2 gene

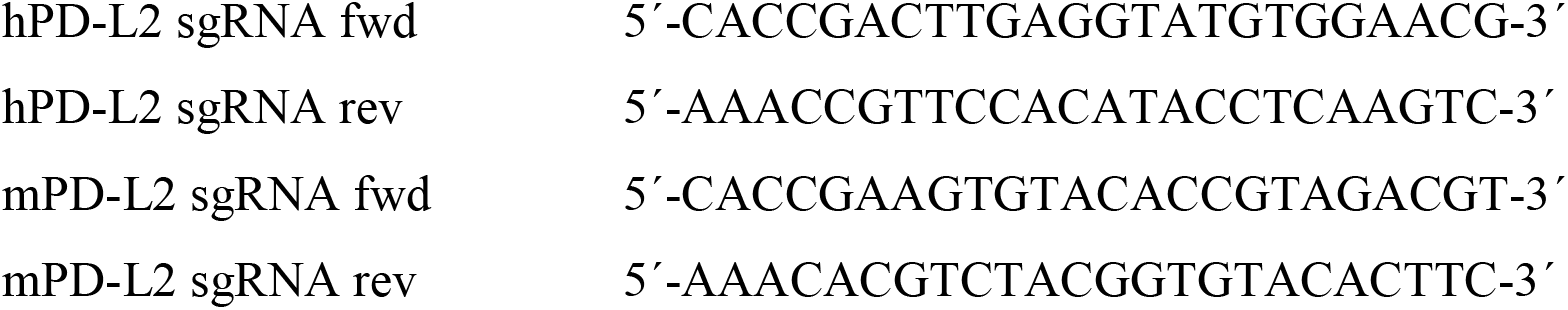

### PCR of genomic DNA

Genomic DNA was isolated using the DNeasy blood and tissue kit (Qiagen) following manufacturer’s instructions. 500 ng of genomic DNA were added to a mix with 0.2 mM dNTP, 1 μM of primers hybridizing to intronic regions spanning exon 3 of PD-L2 (Table 3) and 1 U of BIOTAQ DNA polymerase (Ecogen). After 35 cycles of amplification, the PCR products were separated by electrophoresis on a 1% agarose gel. PCR bands were gel purified using QIAquick gel extraction kit (Qiagen). Isolated PCR fragments were then sent for sequencing to Eurofins Genomics and analysed using Serial Cloner V2.6.1.

Primers used for PCR of exon 3 of human and mouse PD-L2 gene

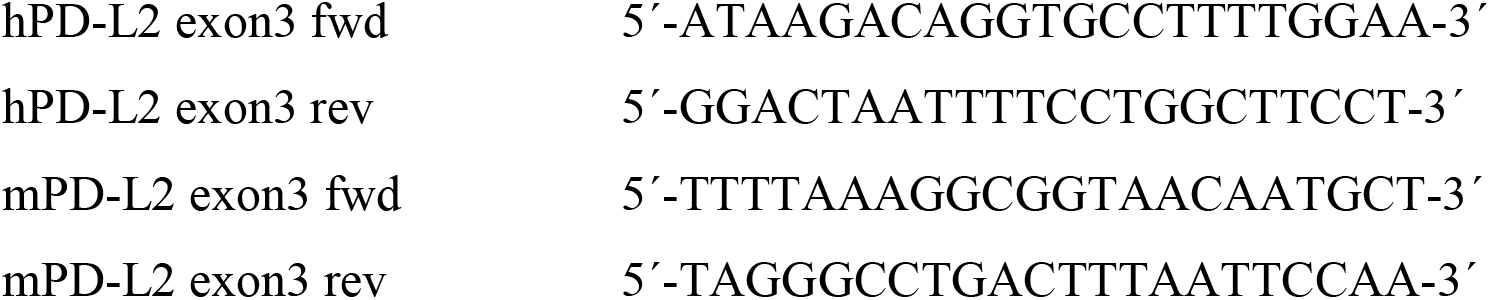

### Transfections

Cells were transfected with pSpCas9(BB)-2A-Puro (PX459) (Addgene #48139) using FuGENE6 (Promega) following the manufacturer’s recommendations. A total of 3 μg of sgRNA containing PX459 plasmid were transfected. After three days, successfully transfected cells were selected using puromycin (Merck). Cells were then either used in bulk or single clones were isolated by plating 0.5 cells per well in a 96 well plate until colony formation. Successfully generated knockouts from single cell colonies were then assessed by sequencing of sgRNA targeted exons, IHC and flow cytometry for genome editing and PD-L2 protein expression respectively.

### Generation of lentiviruses and infections

Lentiviruses were produced by transfecting HEK293T cells with p8.91 (gag-pol expressor), pMDG.2 (VSV-G expressor) and lentiCRISPRv2 (Addgene #52961) for generation of bulk PD-L2-KO cells or luciferase (Addgene #105621) to monitor in vivo bioluminescence following standard procedures. Virus batches were harvested 48, 72 and 96 hours after transfection. Cellular debris was removed by centrifugation and filtering. The supernatant containing the virus was then used fresh or stored by snap-freezing aliquots supplemented with 8 μg/ml polybrene (Fisher Scientific) in ethanol/dry ice and stored at -80° C. Recipient cells were incubated for eight hours with lentivirus and selected three days later with puromycin (Merck) or G418 (Thermofisher) for lentiCRISPRv2 or luciferase experiments, respectively.

### Plasma membrane proteomic screening

SK-MEL-103 cells were induced to senesce by addition of 200 nM doxorubicin for 48 h, or 1 µM palbociclib for the duration of the experiment. At day 7, up to 5 × 10^6^ cells per condition, including growing cells seeded 48 h before as controls, were collected in cold PBS by scraping and pelleted by centrifugation. Plasma membrane proteins were extracted using a plasma membrane protein extraction kit (Abcam, #ab65400), following the manufacturer’s instructions. Proteins were dissolved in UT buffer (8 M urea, 2 M thiourea, 100 mM Tris-HCl pH = 8.0) and digested using a standard FASP protocol. The proteins were reduced (15 mM TCEP, 30 minutes, room temperature, RT), alkylated (50 mM CAA, 20 minutes in the dark, RT) and sequentially digested with Lys-C (Wako) (protein:enzyme ratio 1:50, overnight at RT) and trypsin (Promega) (protein:enzyme ratio 1:100, 6 h at 37° C). Resulting peptides were desalted using Sep-Pak C18 cartridges (Waters). LC-MS/MS was performed by coupling an UltiMate 3000 RSLCnano LC system to either a Q Exactive HF or Q Exactive HF-X-mass spectrometer (Thermo Fisher Scientific). In both cases, peptides were loaded into a trap column (Acclaim™ PepMap™ 100 C18 LC Columns 5 µm, 20 mm length) for 3 minutes at a flow rate of 10 µl/minute in 0.1% FA. Then, peptides were transferred to an EASY-Spray PepMap RSLC C18 column (Thermo) (2 µm, 75 µm x 50 cm) operated at 45° C and separated using a 90 min effective gradient (buffer A: 0.1% FA; buffer B: 100% ACN, 0.1% FA) at a flow rate of 250 nl/min. The gradient used was: from 4% to 6% of buffer B in 2.5 min, from 6% to 25% B in 72.5 min, from 25% to 42.5% B in 14 min plus 6 additional min at 98% B. Peptides were sprayed at 1.8 kV into the mass spectrometer via the EASY-Spray source and the capillary temperature was set to 300° C. The Q Exactive HF was operated in a data-dependent mode, with an automatic switch between MS and MS/MS scans using a top 15 method (intensity threshold ≥ 6.7e4, dynamic exclusion of 26.25 sec and excluding charges +1 and > +6). MS spectra were acquired from 350 to 1400 m/z with a resolution of 60,000 FWHM (200 m/z). Ion peptides were isolated using a 2.0 Th window and fragmented using higher-energy collisional dissociation (HCD) with a normalised collision energy of 27. MS/MS spectra resolution was set to 15,000 or 30,000 (200 m/z). The ion target values were 3e6 for MS (maximum IT of 25 ms) and 1e5 for MS/MS (maximum IT of 15 or 45 msec). The Q Exactive HF-X was operated in a data-dependent mode, with an automatic switch between MS and MS/MS scans using a top 12 method (intensity threshold ≥ 3.6e5, dynamic exclusion of 34 sec and excluding charges +1 and > +6). MS spectra were acquired from 350 to 1400 m/z with a resolution of 60,000 FWHM (200 m/z). Ion peptides were isolated using a 1.6 Th window and fragmented using higher-energy collisional dissociation (HCD) with a normalised collision energy of 27. MS/MS spectra resolution was set to 15,000 (200 m/z). The ion target values were 3e6 for MS (maximum IT of 25 ms) and 1×10^5^ for MS/MS (maximum IT of 22 ms). Raw files were processed with MaxQuant using the standard settings against either a human protein database (UniProtKB/Swiss-Prot, 20373 sequences) or a mouse database (UniProtKB/TrEMBL, 53449 sequences). Carbamidomethylation of cysteines was set as a fixed modification whereas oxidation of methionines and protein N-term acetylation were set as variable modifications. Minimal peptide length was set to 7 amino acids and a maximum of two tryptic missed-cleavages were allowed. Results were filtered at 0.01 FDR (peptide and protein level). Afterwards, the “proteinGroups.txt” file was loaded in Prostar (Wieczorek et al, Bioinformatics 2017) using the LFQ intensity values for further statistical analysis. Briefly, proteins with less than 75% valid values in at least one experimental condition were filtered out. When needed, a global normalisation of log2-transformed intensities across samples was performed using the LOESS function. Missing values were imputed using the algorithms SLSA^56^ for partially observed values and DetQuantile for values missing on an entire condition. Differential analysis was performed using the empirical Bayes statistics Limma. Proteins with a p-value < 0.05 and a log_2_ ratio >0.58 (1.5 in non-log scale) were defined as upregulated. The FDR was estimated to be below 5%.

### Transcriptomic analysis and gene set enrichment analysis

RNA was extracted from PD-L2-WT and PD-L2-KO Panc02 cells, treated with 200 nM doxorubicin for 48 h or untreated, using the RNeasy Mini Kit (Qiagen). The concentration of total RNA was quantified with the Nanodrop One (Thermo Fisher) and RNA integrity was assessed with the Bioanalyzer 2100 RNA Nano assay (Agilent). Libraries for RNA-seq were prepared at the IRB Barcelona Functional Genomics Core Facility. Briefly, mRNA was isolated from 1.5 μg of total RNA using the kit NEBNext Poly(A) mRNA Magnetic Isolation Module (New England Biolabs). NGS libraries were prepared from the purified mRNA using the NEBNext Ultra II RNA Library Prep Kit for Illumina (New England Biolabs). Ten cycles of PCR amplification were applied to all libraries. The final libraries were quantified using the Qubit dsDNA HS assay (Invitrogen) and quality controlled with the Bioanalyzer 2100 DNA HS assay (Agilent). An equimolar pool was prepared with the sixteen libraries and sequenced on a NextSeq550 (Illumina) at IRB. 42.5 Gbp of SE75 reads were produced from a High Output run. A minimum of 32.2 million reads were obtained for all samples. Single-end reads were aligned to the mouse genome version mm10 using STAR (v.2.5.2b). SAM files were converted to BAM files and sorted using sambamba. The count matrix was generated with Rsubread^57^ with the built-in annotation for mm10. DESEq2^58^ was used for differential expression analysis with fold change shrinkage as implemented in the lfcShrink function. Functional enrichment analysis was performed over gene sets defined in the Molecular Signatures Database (MSigDB) hallmark gene set collection. The rotation-based approach for enrichment^59^ implemented in the R package limma was used to represent the null distribution. The max-mean enrichment statistic proposed elsewhere, under restandardisation, was considered for competitive testing.

### Drug class enrichment analysis

Datasets from the LINCS L1000 project (http://www.lincsproject.org/) were analysed and differential expression of *PDCD1LG2* was detected by comparing control and treated samples introducing a correction by cell line and batch effect^60^. 4690 drugs with common names were then further processed and categorised into drugs sets reflecting their mode of action. The resulting drug set enrichment analysis was then compared to a drugset consisting of random drugs.

### Mouse husbandry

All procedures were approved by the Ethical Committee for Animal Experimentation (CEEA) at the Parc Cientific de Barcelona (Spain), the CNIO-ISCIII Ethics Committee for Research and Animal Welfare (Madrid, Spain) and the Ethical Committee for Animal Experimentation (CEEA) at Vall d’Hebron Institut de Recerca (VHIR, Barcelona, Spain). Male wildtype mice (C57BL/6) were purchased from Charles River. Male nude mice (Hsd:athymic nude-*Foxn1*^*nu*^) were purchased from Envigo. MMTV-PyMT (PyMT) mice, in a FVB background, were purchased from The Jackson Laboratory (Bar Harbor, ME). The animals were kept under a 12 h-12 h light-dark cycle and allowed unrestricted access to food and water.

### Syngeneic Panc02 pancreatic tumours

Syngeneic pancreatic adenocarcinoma Panc02 cells expressing firefly luciferase were orthotopically injected in the pancreas of immunocompetent or nude mice (5 × 10^5^ cells in 50 µl PBS). Tumour growth was assessed once a week using an IVIS Spectrum Imaging System (Perkin

Elmer Inc). 10 minutes after intraperitoneal injection of 75 mg/kg luciferin, bioluminescence was recorded. Quantification of tumour burden was performed using Living Image 3.2 software (Perkin Elmer Inc). Doxorubicin (Sigma-Aldrich) treatment (or PBS as vehicle) was applied in the indicated experimental groups at days 7, 10 and, unless otherwise noted, 24 after surgery. Survival was monitored until the animals reached the humane endpoints related to tumour size, anaemia, subcutaneous oedema or ascites. Doxorubicin was injected at 4 mg/kg, intraperitoneally. For short-term determinations, additional cohorts of mice were euthanised at day 12, and the tumours were formalin-fixed for immunohistochemistry or processed for mass cytometry.

### Treatments with blocking antibodies

For selective elimination of immune cell populations in C57BL/6 mice with Panc02 orthotopic tumours, the animals were treated twice a week, starting three days after the surgery, with 100 µg each of, alternatively, anti-CD4 (clone GK1.5, BioXCell #BP0003-1), anti-CD8α (clone 2.43, BioXCell #BE0061) or IgG2b (clone LTF-2, BioxCell #BP0090) as an isotype control. Doxorubicin was administered on days 7 and 10 as described. Tumour growth was monitored by IVIS and the mice were euthanised at day 28. Alternatively, mice with Panc02 orthotopic tumours were treated twice a week, starting three days after the surgery, with 200 µg anti-Gr1 (clone RB6-8C5, BioXCell #BE0075) or isotype control (clone LTF-2, BioXCell #BP0090). Doxorubicin was administered as described on days 7, 10 and 24 and the mice were euthanised at day 35.

### Determination of circulating immune populations by flow cytometry

Approximately 200 µl of total blood of mice treated with doxorubicin and anti-CD4/8 blocking antibodies was extracted by terminal cardiac puncture and kept briefly on ice in a tube containing EDTA. 10 ml of red blood cells (RBC) lysis buffer (BioLegend) were added and the samples were incubated for 5 min at 37º C. The samples were then centrifuged at 350 x g and washed with PBS twice and in 100 µl FACS buffer once, before an additional centrifugation and resuspension in 1:400 CD16/CD32 (BD Fc BlockTM, BD Biosciences #553142) for 15 minutes, at 4º C. 50 µl of a primary antibody mix were later added, containing 1:500 anti-CD45-ApCCy7 (Biolegend #103116), 1:300 anti-CD3-APC (Thermo Fisher #17-0032-80), 1:400 anti-CD4-PerCP Cy5.5, (BD Pharmingen #550954), 1:200 anti-CD8-FITC, (Thermo Fisher #11-0081-82) for an incubation of 25 minutes at 4º C. After the incubations, the samples were washed three times with FACS buffer, resuspended in 250 µl FACS buffer with 1 µl DAPI and analyzed with a Gallios flow cytometer (Beckman Coulter)

### Other syngeneic tumour mouse models

0.4 × 10^6^ HCmel3 cells were subcutaneously injected into male C57BL/6 mice. Four weeks later mice were treated bi-weekly with 5 mg/kg intravenous (i. v.) doxorubicin for a total of three times. 0.2 × 10^6^ B16OVA WT or B16OVA PD-L2KO cells were subcutaneously injected into male C57BL/6 mice. Mice were treated then on day 7 and day 10 with 5 mg/kg doxorubicin (i. v.) and on day 17 with 5 mg/kg intraperitoneal doxorubicin. Tumour growth was monitored by caliper measurements and tumour volume calculated using the formula volume = (length * width^2^) / 2. Mice were euthanised two days after the last treatment.

### SK-MEL-103 xenograft tumours

10^6^ SK-MEL-103 cells, either PD-L2-WT or PD-L2-KO, were harvested, resuspended in a volume of 100 µl PBS and injected in the flank of athymic nude mice. Once tumours the were visible, approximately at day 8-10, the mice were randomly assigned to the palbociclib-treated group, which received 100 mg/kg of palbociclib by oral gavage in 50 mM sodium lactate every day, or the control group, which received vehicle. Tumour size was monitored for ten days using a caliper as described above. The tumours were then extracted and flash frozen for further analysis of gene expression or whole mount SABG staining, and formalin fixed for PD-L2 staining as described above.

### Immunohistochemistry in Panc02 pancreatic tumours

Samples were fixed overnight at 4º C with neutral buffered formalin (Sigma-Aldrich). Paraffin-embedded tissue sections (2-3 μm in thickness) were air dried and further dried at 60º C overnight. For p21 detection, immunohistochemistry (IHC) was performed using a Ventana discovery XT for p21 clone HUGO 291H/B5 (CNIO), undiluted, for 60 min. Antigen retrieval was performed with Cell Conditioning 1 (CC1) buffer (Roche, #950-124). Secondary antibodies used were the OmniMap anti-rat HRP (Roche, #760-4457). For Ly6G/C (Gr1) staining, IHC was performed using a Ventana discovery XT for Ly6G/C (Novus #NB600-1387) primary antibody at 1:100 for 60 min. Antigen retrieval was performed with Cell Conditioning 1 (CC1) buffer (Roche, 950-124). After incubation with the primary antibody, the rabbit anti-rat IgG (AI-4001, Vector) at 1:500 for 32 min was used followed with the secondary antibody OmniMap anti-rabbit HRP (760-4311, Roche). Antigen-antibody complexes were revealed with ChromoMap DAB Kit (Roche, 760-159). Specificity of staining was confirmed with the rat IgG isotype control (R&D Systems, #6-001-F). Brightfield images were acquired with a NanoZoomer-2.0 HT C9600 digital scanner (Hamamatsu) equipped with a 20X objective. All images were visualised with a gamma correction set at 1.8 in the image control panel of the NDP.view 2 U12388-01 software (Hamamatsu, Photonics, France). Automated quantification was performed using QuPath^61^.

### Senescence-associated β-galactosidase stainings

SABG stainings of adherent cells, OCT embedded tumours or whole mount tumours were performed using homemade solutions adapted from the original protocol^62^. A fixation solution was prepared containing 5 mM EGTA, 2 mM MgCl_2_ and 0.2% glutaraldehyde in 0.1 M phosphate buffer (pH 7). A staining solution was prepared containing 40 mM citric acid, 5 mM potassium cynoferrate (II), 5 mM potassium cyanoferrate (III), 150 mM sodium chloride and 2 mM magnesium chloride in 0.1 M phosphate buffer (pH 6). The solution was light protected and stored at 4° C. Prior to staining, the needed amount of staining solution was pre-warmed at 37° C and 5-bromo-4-chloro-3-indolyl ß-D-galactoside (X-Gal) was added to a final concentration of 1 mg/ml. X-gal was added directly before the staining from a 100 mg/ml solution in N,N dimethylformamide, stored at -20° C for no longer than 4 weeks. For staining of cells, they were washed with PBS, an appropriate amount of cold fixation solution was added and cells were incubated for 15 minutes at RT. After that, cells were washed twice with PBS and then the X-gal-containing staining solution was added. Cells were incubated in a non-CO_2_ regulated incubator at 37° C overnight. The following day, the staining solution was removed, cells were washed twice with PBS and stored in 50% glycerol at 4° C until the analysis was performed. For whole tumour SABG staining, the staining procedure is the same as for cells with the modification that tissues were fixed for 45 minutes in fixation solution at RT and then incubated with staining solution for 16 hours at 37 °C. For OCT-embedded, frozen tumours, 4 µm-thick sections were obtained, followed by fixing and staining as described above. Nuclear fast red was used as counterstain.

### Identification of immune cell populations by mass cytometry (CyTOF)

Single cell suspensions were prepared from Panc02 pancreatic tumours. The tumours were briefly kept in DMEM on ice and manually minced in DMEM. The tumour fragments were incubated in 10 mg/ml collagenase I and DNase in DMEM for 30 minutes, at 37º C, with gentle shaking, in gentleMACS C tubes, with several steps of processing in a gentleMACS Dissociator before and after the incubation following the manufacturer’s instructions. The resulting cell suspension was passed through a 70 µm strainer, washed in PBS, incubated in RBC lysis buffer (BioLegend) for 5 minutes at room temperature and then washed with PBS a second time. The cells were then resuspended incubated with anti-mouse CD16/CD32 at 1:400 (BD Fc BlockTM, BD Biosciences #553142) for 10 minutes, at 4º C in 50 µl in Maxpar Cell Staining Buffer (CSB). Then, an equal volume of the 16-antibody cocktail from the Maxpar mouse spleen/lymph node phenotyping panel kit (Fluidgm #201306), spanning all major immune cell populations, was added and incubated for 15 minutes at room temperature, vortexed and incubated for 15 additional minutes. The cell suspension was then washed twice with 2 ml Maxpar CSB and fixed in 1.6% formaldehyde in PBS for 10 min at room temperature. The samples were then centrifuged at 800 x g for 5 minutes and resuspended in 1 ml of 125 nM Cell-ID Intercalator-Ir (Fluidigm) in Maxpar Fix and Perm Buffer (Fluidigm), followed by vortex and incubation 4º C overnight. Acquisition was performed the following day after washing with Cell Acquisition Solution (Fluidigm) and mixing with diluted EQ bead solution (Fluidigm), following the manufacturer’s instructions, in a Helios detector (Fluidigm). The gating strategy is fully standardized and described in detail by the manufacturer. Briefly, Intercalator-Ir positive cells are selected, from which CD19^+^B220^+^ B lymphocytes are defined. Next, TCRβ^+^CD3^+^ T cells are selected and, within this population, CD4^+^ and CD8^+^ T cells are detected. From the TCRβ^-^CD3^-^ population, sequentially, we identified by sequential exclusion NK1.1^+^ NK cells and CD11c^+^ dendritic cells. Finally, among CD11b^+^ cells, macrophages and MDSCs were identified depending on Gr1 positivity.

### Measurement of intratumoural cytokine and chemokine levels

Panc02 pancreatic tumours were flash frozen after excision and approximately 10 mg of tumour mass was homogeneised in RIPA lysis buffer (Merck) using a FastPrep-24 5G homogeneiser (MP Biomedicals) in the presence of Halt protease inhibitor cocktail (Thermo Scientific). The protein concentration was determined by a colorimetric assay (DC protein assay kit, BioRad). All samples were diluted to a protein concentration of 1.2 mg/ml in a total volume of 75 µl of RIPA and shipped to Eve Technologies (Calgary, Canada) on dry ice. The intratumoural levels of CCL11 (eotaxin), G-CSF, GM-CSF, IFN-γ, IL-1α, IL-1β, IL-2, IL-3, IL-4, IL-5, IL-6, IL-7, IL-9, IL-10, IL-12p40, IL-12p70, IL-13, IL-15, IL-17, CXCL10 (IP 10), CXCL1 (KC), LIF, CXCL6 (LIX), CCL2 (MCP-1), CXCL9 (MIG), CCL3 (MIP-1α), CCL4 (MIP-1β), CXCL2 (MIP-2), CCL5 (RANTES), TNFα and VEGF were determined using a Multiplexing LASER Bead Assay (Mouse Cytokine Array / Chemokine Array 31-Plex, Eve Technologies). For the cytokines and chemokines included in the assay but not shown in the manuscript (IL-13, IL-12p40, CCL4), no changes were observed.

### MMTV-PyMT model

Male PyMT mice were mated with FVB-WT female to generate female littermates that are PyMT/wild-type (WT). Mice were genotyped for the PyMT allele by polymerase chain reaction (PCR) using the following primers: forward primer, 5’-CGCACATACTGCTGGAAGAA and reverse primer, 5’-TGCCGGGAACGTTTTATTAG. Weights were recorded for every experiment. The mice were housed in a specific pathogen–free environment. At 9 weeks of age, PyMT mice were palpated to detect the onset of mammary tumour development. The mice were then injected intraperitoneally with 4 mg/kg doxorubicin or an equivalent volume of PBS, once a week for 4 weeks. Additional groups of mice were treated with anti-mouse PD-L2 (clone TY25, BioXCell, #BE0112), alone or in combination with doxorubicin. From the detection of palpable tumours, the mice were monitored for tumour growth with a caliper twice a week. Total tumour volume was determined using the formula: volume = (length * width^2^) / 2. The mice were euthanised 4 weeks after the onset of tumours and the start of the treatments. For depletion of specific immune populations, the mice were injected intraperitoneally with anti-CD4 (clone YTS 191, BioXCell #BE0119), anti-CD8α (clone 53-6.7, BioXCell #BE0004-1), or rat IgG (IgG2a isotype, BioXCell #BE0089) and euthanized at week 13. All the antibody treatments were administered i. p., every three days for four weeks, at a 10 mg/kg dose.

### Immunohistochemistry and SABG in mammary tumours

For PMyT mice, mammary glands were either formalin-fixed or whole mounted, as previous described, and paraffin-embedded. IHC experiments were performed using the Dako EnVision+ System-HRP kit. Using the Leica microtome system, 3 μm-thick sections were obtained, and antigen retrieval was performed using Tris EDTA buffer (pH 9) by boiling followed by washing in running water and blocking for endogenous peroxidase for 15 min at room temperature. The slides were washed with Tris-buffered saline (TBS) and, after 1 h of incubation with 3% BSA blocking solution, each of them was incubated with primary antibody against PD-L2 (1:100, CST #49189), CD3 (1:100, Abcam #ab16669), CD4 (1:1000, Abcam #ab183685) or CD8 (Abcam #1:2000, ab217344) at 4° C overnight. Followed by washing, they were incubated with System-HRP Labelled Polymer Anti-Rabbit (Dako, K400311-2) secondary antibodies for 1 hour at room temperature. The tissues were counterstained with hematoxylin after the substrate (DAB) reaction step. For SABG staining, mammary gland samples were whole mounted using a fixative solution of 0.2% glutaraldehyde, 2% paraformaldehyde in PBS. Overnight staining with senescence β-galactosidase staining kit (CST) was performed directly in the fixed samples, previous to their inclusion in paraffin. Using the Leica microtome system, 3 μm-thick sections were obtained and counterstained with Liquid-Stable Nuclear Fast Red (VWR Life Science AMRESCO).

### Quantification and Statistical Analysis

Results are presented as mean ± SEM for all data. All statistical analysis were performed using GraphPad Prism 8 and a p value lower than 0.05 was considered significant. Tests were applied as described in the figures, with the general use of two-tailed Student’s t-tests for pairwise comparisons between two isolated conditions and one-way ANOVA with Tukey’s multiple comparison for experiments with three or more experimental conditions. For datasets including two independent factors, in which one of them is usually time, 2-way ANOVA was used. Specific tests for select experiments are detailed in the figure legends and/or in the methodological description.

## Data availability

The proteomic screen data is available at ProteomeXchange located at http://proteomecentral.proteomexchange.org/cgi/GetDataset (Project accession: PXD033714; Username; Password: EkVHvHFo). The RNA sequencing data has been submitted to Gene Expression Omnibus (GEO) and are pending to obtain an accession number.

## Acknowledgements

J.A.L-D. was supported by the Spanish Ministry of Science through a Juan de la Cierva-Incorporación fellowship and by the Spanish Association Against Cancer (AECC) through an AECC Investigador fellowship. I.M. was funded by a FPI fellowship from the Spanish Ministry of Science. Work in the laboratory of M.S. was funded by the IRB and “laCaixa” Foundation, and by grants from the Spanish Ministry of Science co-funded by the European Regional Development Fund (ERDF) (SAF2017-82613-R), European Research Council (ERC-2014-AdG/669622), and Secretaria d’Universitats i Recerca del Departament d’Empresa i Coneixement of Catalonia (Grup de Recerca consolidat 2017 SGR 282). J.L.K., T.T., and S.C. are supported by the National Institutes of Health (grants R37AG13925, R33AG61456, R01AG072301, R01AG61414, P01AG62413, and UH3AG56933), Robert and Arlene Kogod, the Connor Fund, Robert J. and Theresa W. Ryan, and the Noaber Foundation. Work in the laboratory of J.A. is supported by the Breast Cancer Research Foundation (BCRF-21-008), Instituto de Salud Carlos III (project references AC15/00062, CB16/12/00449 and PI19/01181) and the EC (under the Framework of the ERA-NET TRANSCAN-2 initiative co-financed by FEDER), AECC and Fundació La Caixa (HR22-00776).

## Author contributions

S.C., J.A.L.-D. and M.L. designed and performed experiments and cowrote the manuscript. M.A. and N.P. performed histopathological analyses. I.M., K.M., S.L. and F.A.-S. provided data and feedback. M.M.M, S.P.-E. and M.E. provided technical support. L.M. and C.S.-O.A. performed computational analysis. T.C., T.T. and J.L.K. provided discussion and revisions. J.A. and M.S. designed and supervised the study, secured funding analysed the data and cowrote the manuscript.

## Additional information

Extended Data is available for this paper. Correspondence and requests for materials should be addressed to Manuel Serrano.

---

**Extended Data Fig. 1.**
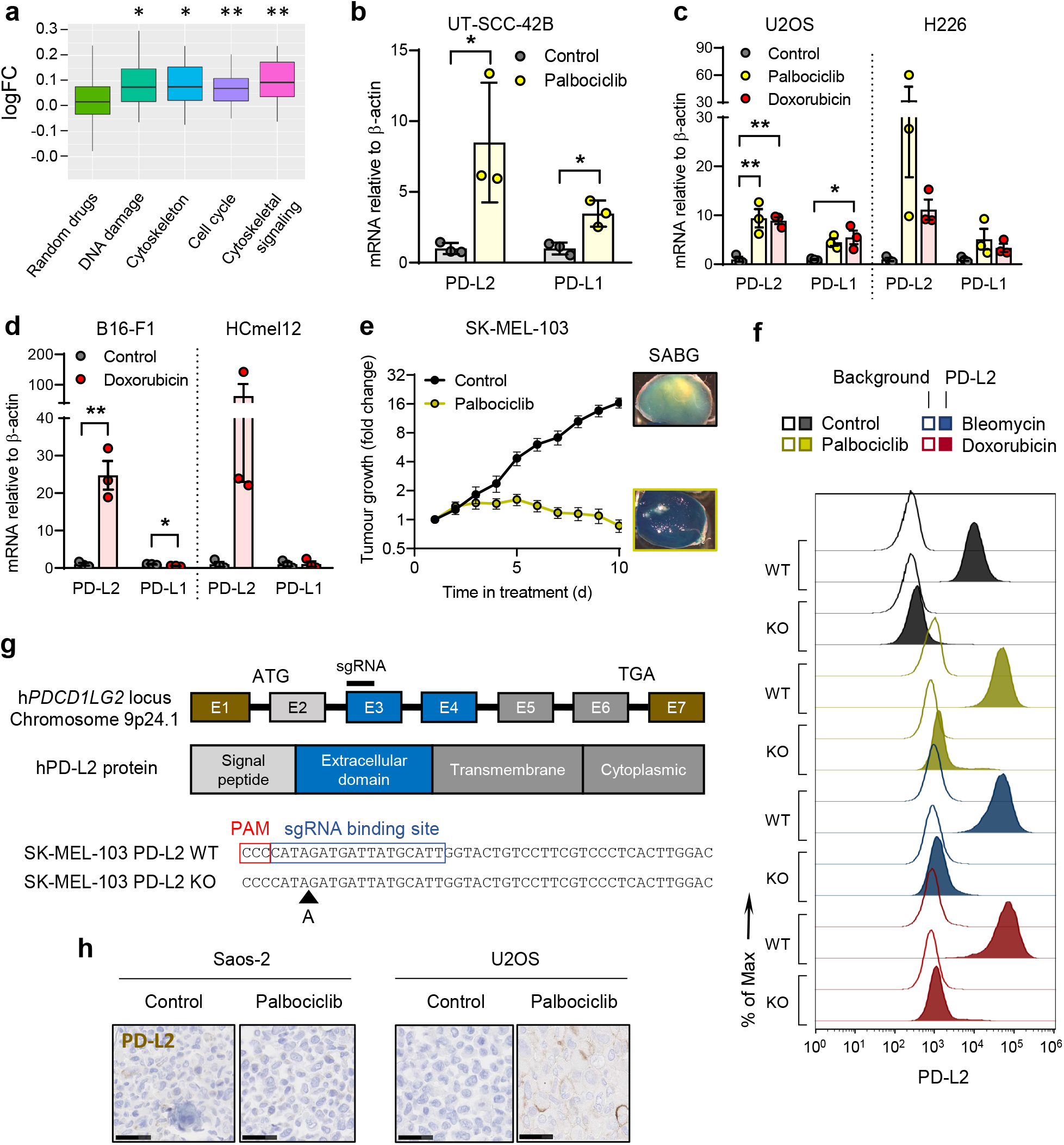
PD-L2 is upregulated in human and murine senescent cancer cells. (a) Drug class enrichment analysis for human *PDCD1LG2* (PD-L2). (b) PD-L1/2 mRNA expression in human cancer cell lines treated with palbociclib. (c) PD-L1/2 mRNA expression in human cancer cell lines after treatment with doxorubicin or palbociclib. (d) PD-L1/2 mRNA expression in murine cancer cell lines treated with doxorubicin. (e) Growth chart of SK-MEL-103 xenograft tumours in nude mice, untreated or treated with palbociclib, and SABG staining in toto of representative tumours. (f) Representative example (1 out of n =3) of histogram for PD-L2 protein levels upon generation of a PD-L2-KO SK-MEL-103 cell line, in control and senescent conditions, measured by flow cytometry. (g) CRISPR-Cas9 edition of the human *PDCD1LG2* locus, specifying the sgRNA biding site in exon 3. The sequence corresponds the single clone of edited SK-MEL-103 cells used in the experiments labelled as PD-L2-KO SK-MEL-103. (h) PD-L2 staining in Saos-2 and U2OS cell pellets, treated with palbociclib. Scale bars = 50 µm. t-tests or 1 way ANOVA with Tukey post-test were applied. * p < 0.05, ** p < 0.01, *** p < 0.001.

**Extended Data Fig. 2.**
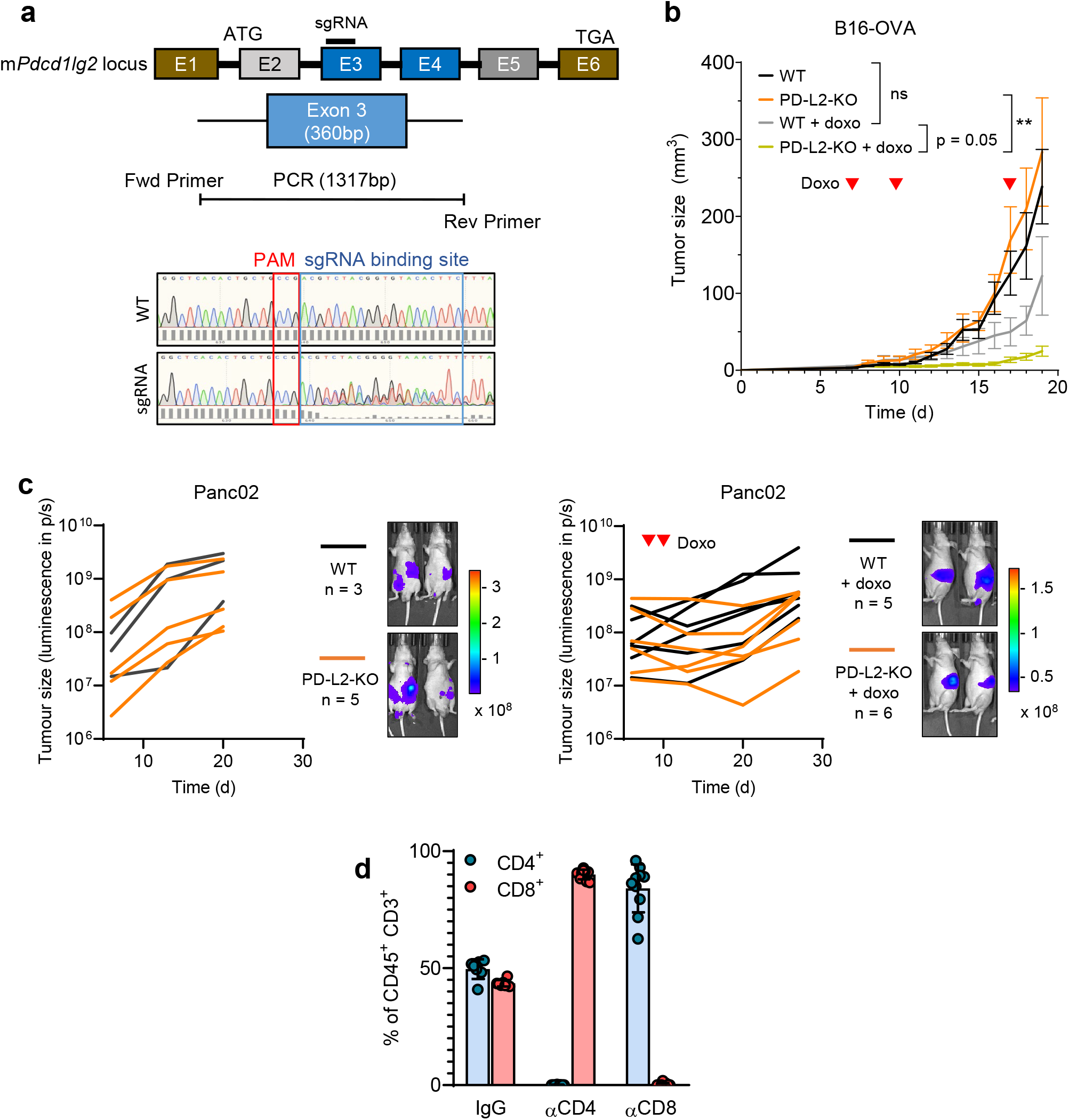
A combination of PD-L2 ablation and chemotherapy results in tumour remission. (a) CRISPR edition of the murine *Pdcd1lg2* locus, specifying the sgRNA biding site in exon 3, that generated a bulk population of edited Panc02 cells. This bulk population was used in all the experiments labelled as PD-L2-KO Panc02. (b) Growth chart of WT and bulk PD-L2-KO B16-OVA tumours in C57BL/6 mice, untreated or treated with doxorubicin on days 7, 10 and 17 after subcutaneous injection of cells. B16-OVA-WT n = 5, B16-OVA-KO n = 6, B16-OVA-WT + doxo n = 11, B16-OVA-KO + doxo n = 11. 2-way ANOVA. (c) Quantification and representative images of Panc02 WT and PD-L2-KO tumours in nude mice, untreated and treated with doxorubicin on days 7 and 10. (d) Percentage of CD4^+^ and CD8^+^ T cells among total CD45^+^ CD3^+^ cells in blood of mice after treatment with blocking anti-CD4 and anti-CD8 antibodies. Luminescence units are photon/second (p/s) in the graphs and photon/sec/cm^2^/stereoradian in the images.

**Extended Data Fig. 3.**
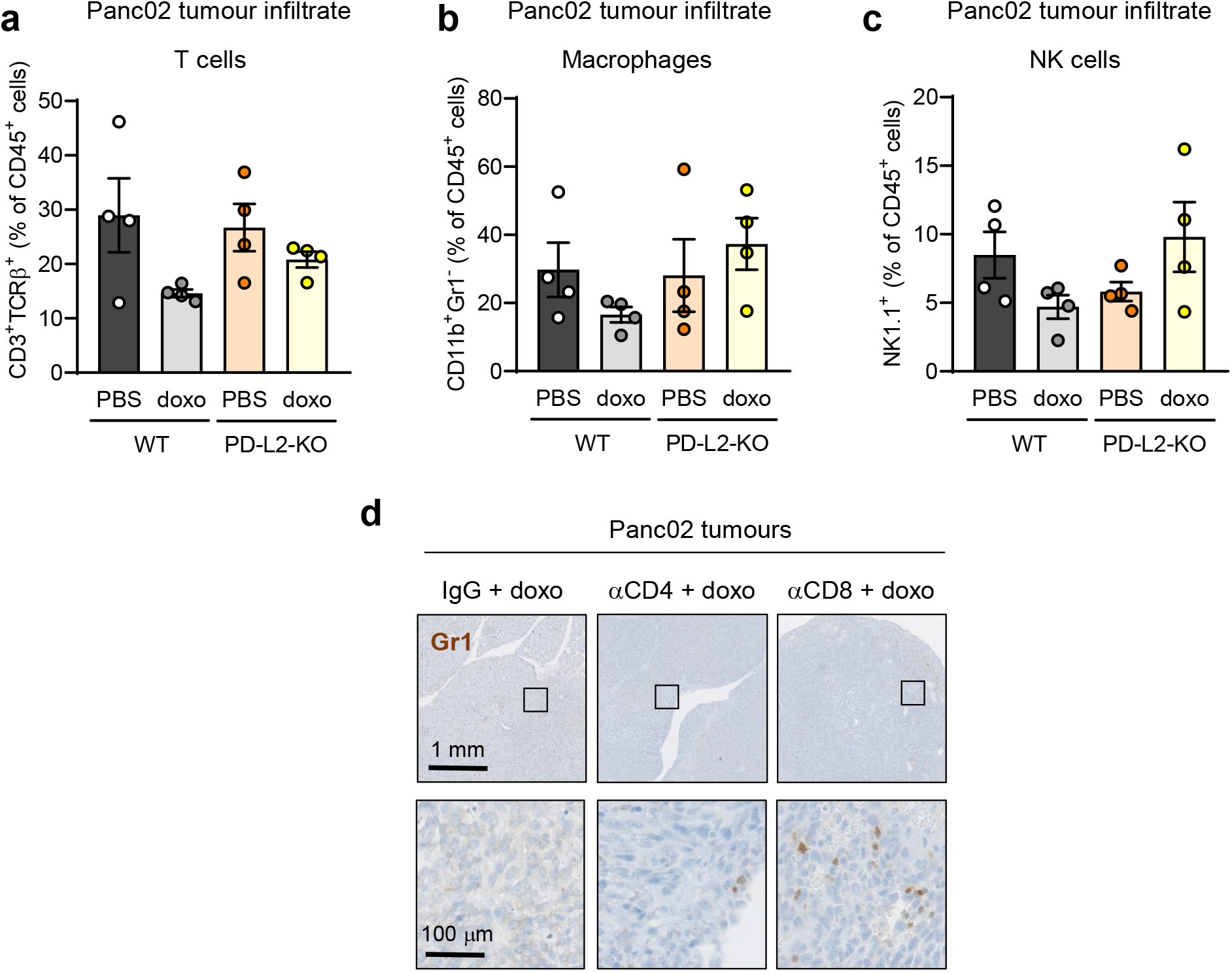
Recruitment of tumour-promoting myeloid cells is prevented in PD-L2-KO tumours upon doxorubicin treatment. (a-c) Quantification of the percentage of (a) lymphocytes (CD3^+^ TCRβ^+^), (b) macrophages (CD11b^+^ Gr1^−^) and (c) NK cells (NK1.1^+^) relative to total CD45^+^ cells, in WT and PD-L2-KO tumours untreated or treated with doxorubicin at days 7 and 10. See methods for further detail in the gating strategies. Samples were collected on day 28. (d) Representative Gr1 stainings in sections of PD-L2-KO tumours treated with doxorubicin and subject to depletion of CD4^+^ or CD8^+^ T cells from Fig. 2c.

**Extended Data Fig. 4.**
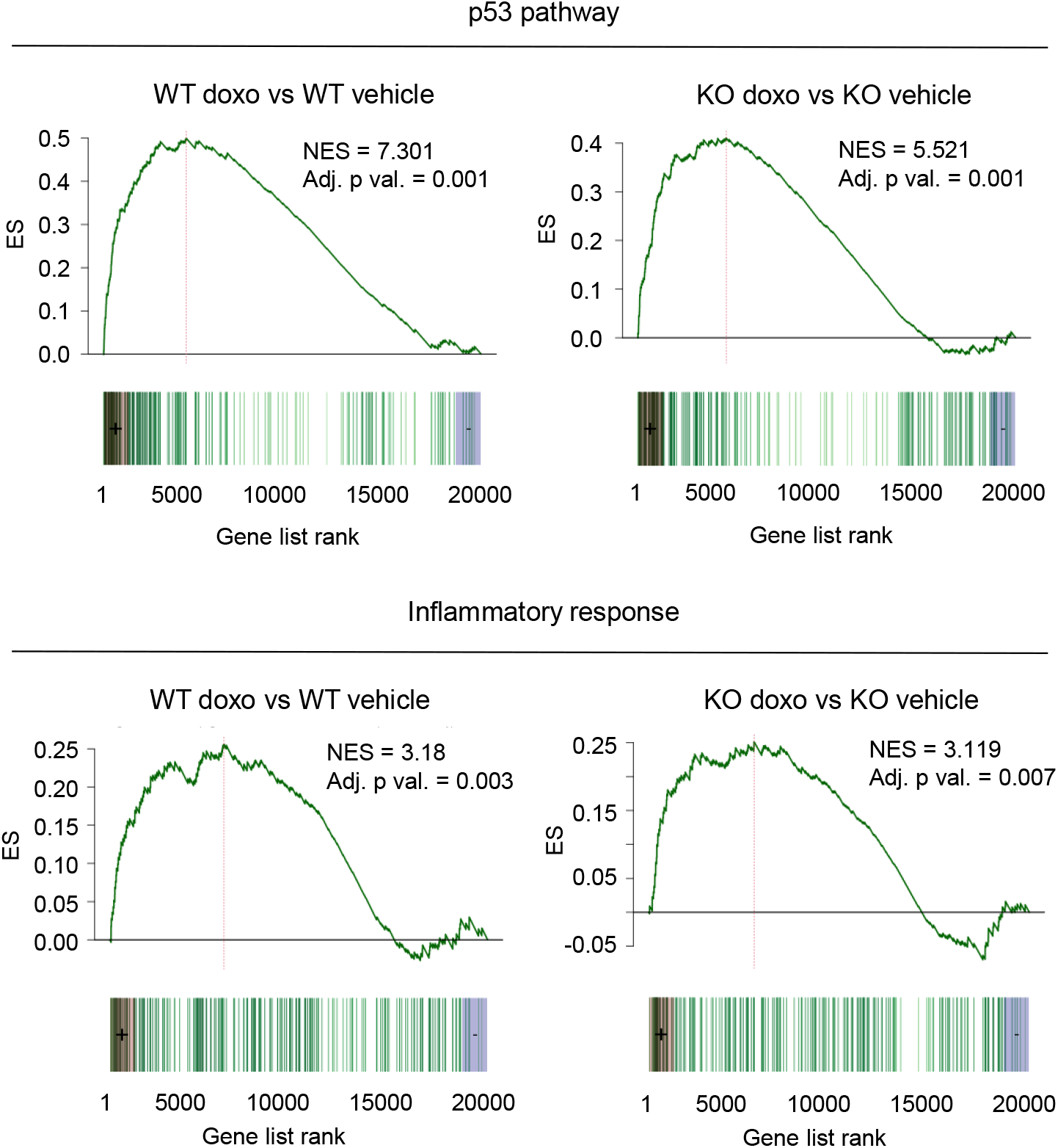
The senescent phenotype of PD-L2-KO Panc02 cells is nearly identical to their WT counterparts. GSEA plots for representative gene sets (Molecular Signatures Database hallmark gene set collection) associated to cellular senescence in doxorubicin-treated versus untreated WT Panc02 cells, as well as in doxorubicin-treated versus untreated PD-L2-KO Panc02 cells at day 7 after treatment.

**Extended Data Fig. 5.**
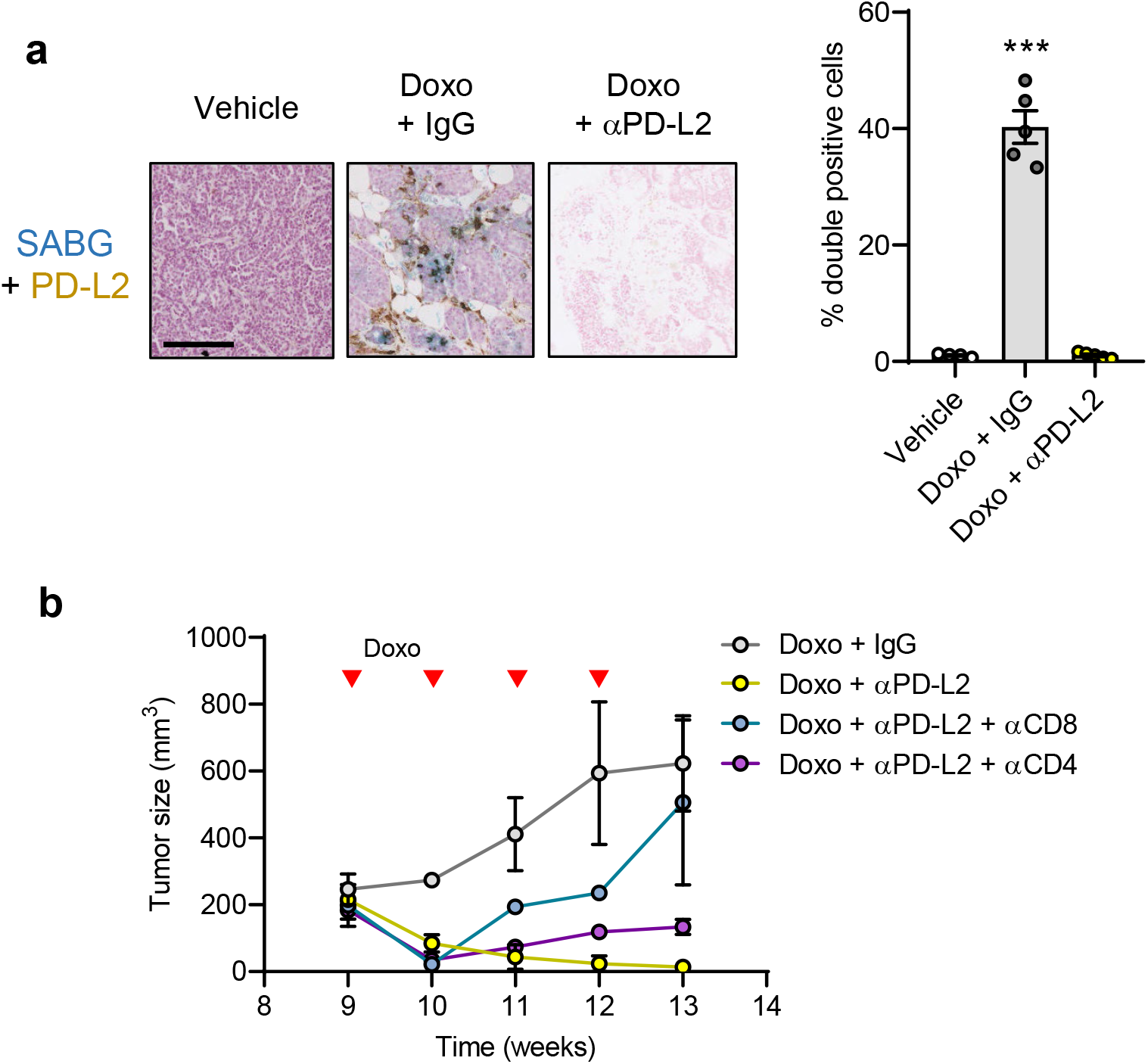
PD-L2 blockade succesfully eliminates mammary tumours after chemotherapy. (a) SABG and PD-L2 costaining in tumours samples from PyMT mice treated weekly with doxorubicin from weeks 9 to 13. Representative images are shown. Samples were obtained at 13 weeks of age, 7 days after the last doxorubicin dose. 1-way ANOVA, *** p < 0.001 (b) Tumour growth in PyMT mice treated with doxorubicin and combinations of blocking antibodies against PD-L2, CD4 and CD8 as indicated. N = 2.

## References

1 Topalian, S. L. et al. Safety, activity, and immune correlates of anti-PD-1 antibody in cancer. N Engl J Med 366, 2443–2454, doi:10.1056/NEJMoa1200690 (2012).

2 Cha, J. H., Chan, L. C., Li, C. W., Hsu, J. L. & Hung, M. C. Mechanisms Controlling PD-L1 Expression in Cancer. Mol Cell 76, 359–370, doi:10.1016/j.molcel.2019.09.030 (2019).

3 Chen, L. & Han, X. Anti-PD-1/PD-L1 therapy of human cancer: past, present, and future. J Clin Invest 125, 3384–3391, doi:10.1172/JCI80011 (2015).

4 Ghosh, C., Luong, G. & Sun, Y. A snapshot of the PD-1/PD-L1 pathway. J Cancer 12, 2735–2746, doi:10.7150/jca.57334 (2021).

5 Ribas, A. & Wolchok, J. D. Cancer immunotherapy using checkpoint blockade. Science 359, 1350–1355, doi:10.1126/science.aar4060 (2018).

6 Sharma, P. & Allison, J. P. The future of immune checkpoint therapy. Science 348, 56–61, doi:10.1126/science.aaa8172 (2015).

7 Solinas, C. et al. Programmed cell death-ligand 2: A neglected but important target in the immune response to cancer? Transl Oncol 13, 100811, doi:10.1016/j.tranon.2020.100811 (2020).

8 Ghiotto, M. et al. PD-L1 and PD-L2 differ in their molecular mechanisms of interaction with PD-1. Int Immunol 22, 651–660, doi:10.1093/intimm/dxq049 (2010).

9 Gorchs, L. et al. Human Pancreatic Carcinoma-Associated Fibroblasts Promote Expression of Co-inhibitory Markers on CD4(+) and CD8(+) T-Cells. Front Immunol 10, 847, doi:10.3389/fimmu.2019.00847 (2019).

10 Lakins, M. A., Ghorani, E., Munir, H., Martins, C. P. & Shields, J. D. Cancer-associated fibroblasts induce antigen-specific deletion of CD8 (+) T Cells to protect tumour cells. Nature communications 9, 948, doi:10.1038/s41467-018-03347-0 (2018).

11 Yearley, J. H. et al. PD-L2 Expression in Human Tumors: Relevance to Anti-PD-1 Therapy in Cancer. Clin Cancer Res 23, 3158–3167, doi:10.1158/1078-0432.CCR-16-1761 (2017).

12 Tanegashima, T. et al. Immune Suppression by PD-L2 against Spontaneous and Treatment-Related Antitumor Immunity. Clin Cancer Res 25, 4808–4819, doi:10.1158/1078-0432.CCR-18-3991 (2019).

13 Latchman, Y. et al. PD-L2 is a second ligand for PD-1 and inhibits T cell activation. Nat Immunol 2, 261–268, doi:10.1038/85330 (2001).

14 Loke, P. & Allison, J. P. PD-L1 and PD-L2 are differentially regulated by Th1 and Th2 cells. Proc Natl Acad Sci U S A 100, 5336–5341, doi:10.1073/pnas.0931259100 (2003).

15 Sudo, S. et al. Cisplatin-induced programmed cell death ligand-2 expression is associated with metastasis ability in oral squamous cell carcinoma. Cancer Sci 111, 1113–1123, doi:10.1111/cas.14336 (2020).

16 Sengedorj, A. et al. The Effect of Hyperthermia and Radiotherapy Sequence on Cancer Cell Death and the Immune Phenotype of Breast Cancer Cells. Cancers (Basel) 14, doi:10.3390/cancers14092050 (2022).

17 Wang, L., Lankhorst, L. & Bernards, R. Exploiting senescence for the treatment of cancer. Nat Rev Cancer, doi:10.1038/s41568-022-00450-9 (2022).

18 Coppe, J. P., Desprez, P. Y., Krtolica, A. & Campisi, J. The senescence-associated secretory phenotype: the dark side of tumor suppression. Annual review of pathology 5, 99–118, doi:10.1146/annurev-pathol-121808-102144 (2010).

19 Krtolica, A., Parrinello, S., Lockett, S., Desprez, P. Y. & Campisi, J. Senescent fibroblasts promote epithelial cell growth and tumorigenesis: a link between cancer and aging. Proc Natl Acad Sci U S A 98, 12072–12077, doi:10.1073/pnas.211053698 (2001).

20 Bavik, C. et al. The gene expression program of prostate fibroblast senescence modulates neoplastic epithelial cell proliferation through paracrine mechanisms. Cancer Res 66, 794–802, doi:10.1158/0008-5472.CAN-05-1716 (2006).

21 Coppe, J. P. et al. A role for fibroblasts in mediating the effects of tobacco-induced epithelial cell growth and invasion. Mol Cancer Res 6, 1085–1098, doi:10.1158/1541-7786.MCR-08-0062 (2008).

22 Birch, J. & Gil, J. Senescence and the SASP: many therapeutic avenues. Genes Dev 34, 1565–1576, doi:10.1101/gad.343129.120 (2020).

23 Demaria, M. et al. Cellular Senescence Promotes Adverse Effects of Chemotherapy and Cancer Relapse. Cancer Discov 7, 165–176, doi:10.1158/2159-8290.CD-16-0241 (2017).

24 Ruhland, M. K. et al. Stromal senescence establishes an immunosuppressive microenvironment that drives tumorigenesis. Nature communications 7, 11762, doi:10.1038/ncomms11762 (2016).

25 Eggert, T. et al. Distinct Functions of Senescence-Associated Immune Responses in Liver Tumor Surveillance and Tumor Progression. Cancer Cell 30, 533–547, doi:10.1016/j.ccell.2016.09.003 (2016).

26 Toso, A. et al. Enhancing chemotherapy efficacy in Pten-deficient prostate tumors by activating the senescence-associated antitumor immunity. Cell reports 9, 75–89, doi:10.1016/j.celrep.2014.08.044 (2014).

27 Salminen, A., Kauppinen, A. & Kaarniranta, K. Myeloid-derived suppressor cells (MDSC): an important partner in cellular/tissue senescence. Biogerontology 19, 325–339, doi:10.1007/s10522-018-9762-8 (2018).

28 Wang, L., Lankhorst, L. & Bernards, R. Exploiting senescence for the treatment of cancer. Nat Rev Cancer 22, 340–355, doi:10.1038/s41568-022-00450-9 (2022).

29 Munoz-Espin, D. et al. A versatile drug delivery system targeting senescent cells. EMBO Mol Med 10, doi:10.15252/emmm.201809355 (2018).

30 Gonzalez-Gualda, E. et al. Galacto-conjugation of Navitoclax as an efficient strategy to increase senolytic specificity and reduce platelet toxicity. Aging Cell 19, e13142, doi:10.1111/acel.13142 (2020).

31 Wyld, L. et al. Senescence and Cancer: A Review of Clinical Implications of Senescence and Senotherapies. Cancers (Basel) 12, doi:10.3390/cancers12082134 (2020).

32 Rivera-Torres, J. & San Jose, E. Src Tyrosine Kinase Inhibitors: New Perspectives on Their Immune, Antiviral, and Senotherapeutic Potential. Front Pharmacol 10, 1011, doi:10.3389/fphar.2019.01011 (2019).

33 Triana-Martinez, F. et al. Identification and characterization of Cardiac Glycosides as senolytic compounds. Nature communications 10, 4731, doi:10.1038/s41467-019-12888-x (2019).

34 Guerrero, A. et al. Cardiac glycosides are broad-spectrum senolytics. Nat Metab 1, 1074–1088, doi:10.1038/s42255-019-0122-z (2019).

35 Wang, C. et al. Inducing and exploiting vulnerabilities for the treatment of liver cancer. Nature 574, 268–272, doi:10.1038/s41586-019-1607-3 (2019).

36 Amor, C. et al. Senolytic CAR T cells reverse senescence-associated pathologies. Nature 583, 127–132, doi:10.1038/s41586-020-2403-9 (2020).

37 Subramanian, A. et al. A Next Generation Connectivity Map: L1000 Platform and the First 1,000,000 Profiles. Cell 171, 1437–1452 e1417, doi:10.1016/j.cell.2017.10.049 (2017).

38 Llanos, S. et al. Lysosomal trapping of palbociclib and its functional implications. Oncogene 38, 3886–3902, doi:10.1038/s41388-019-0695-8 (2019).

39 Krishnamurthy, J. et al. Ink4a/Arf expression is a biomarker of aging. J Clin Invest 114, 1299–1307, doi:10.1172/JCI22475 (2004).

40 Espinosa De Ycaza, A. E. et al. Senescent cells in human adipose tissue: A cross-sectional study. Obesity (Silver Spring) 29, 1320–1327, doi:10.1002/oby.23202 (2021).

41 Xu, M. et al. Targeting senescent cells enhances adipogenesis and metabolic function in old age. Elife 4, e12997, doi:10.7554/eLife.12997 (2015).

42 Schafer, M. J. et al. Cellular senescence mediates fibrotic pulmonary disease. Nature communications 8, 14532, doi:10.1038/ncomms14532 (2017).

43 Jeon, O. H. et al. Local clearance of senescent cells attenuates the development of post-traumatic osteoarthritis and creates a pro-regenerative environment. Nat Med 23, 775–781, doi:10.1038/nm.4324 (2017).

44 Pignolo, R. J., Passos, J. F., Khosla, S., Tchkonia, T. & Kirkland, J. L. Reducing Senescent Cell Burden in Aging and Disease. Trends Mol Med 26, 630–638, doi:10.1016/j.molmed.2020.03.005 (2020).

45 Khosla, S., Farr, J. N., Tchkonia, T. & Kirkland, J. L. The role of cellular senescence in ageing and endocrine disease. Nat Rev Endocrinol 16, 263–275, doi:10.1038/s41574-020-0335-y (2020).

46 Wissler Gerdes, E. O. et al. Cellular senescence in aging and age-related diseases: Implications for neurodegenerative diseases. Int Rev Neurobiol 155, 203–234, doi:10.1016/bs.irn.2020.03.019 (2020).

47 Tchkonia, T. & Kirkland, J. L. Aging, Cell Senescence, and Chronic Disease: Emerging Therapeutic Strategies. JAMA 320, 1319–1320, doi:10.1001/jama.2018.12440 (2018).

48 Farr, J. N. et al. Targeting cellular senescence prevents age-related bone loss in mice. Nat Med 23, 1072–1079, doi:10.1038/nm.4385 (2017).

49 Ovadya, Y. et al. Impaired immune surveillance accelerates accumulation of senescent cells and aging. Nature communications 9, 5435, doi:10.1038/s41467-018-07825-3 (2018).

50 Pereira, B. I. et al. Senescent cells evade immune clearance via HLA-E-mediated NK and CD8(+) T cell inhibition. Nature communications 10, 2387, doi:10.1038/s41467-019-10335-5 (2019).

51 Munoz, D. P. et al. Targetable mechanisms driving immunoevasion of persistent senescent cells link chemotherapy-resistant cancer to aging. JCI Insight 5, doi:10.1172/jci.insight.124716 (2019).

52 Iannello, A., Thompson, T. W., Ardolino, M., Lowe, S. W. & Raulet, D. H. p53-dependent chemokine production by senescent tumor cells supports NKG2D-dependent tumor elimination by natural killer cells. J Exp Med 210, 2057–2069, doi:10.1084/jem.20130783 (2013).

53 Krizhanovsky, V. et al. Senescence of activated stellate cells limits liver fibrosis. Cell 134, 657–667, doi:10.1016/j.cell.2008.06.049 (2008).

54 Sagiv, A. et al. NKG2D ligands mediate immunosurveillance of senescent cells. Aging (Albany NY) 8, 328–344, doi:10.18632/aging.100897 (2016).

55 Prata, L., Ovsyannikova, I. G., Tchkonia, T. & Kirkland, J. L. Senescent cell clearance by the immune system: Emerging therapeutic opportunities. Semin Immunol 40, 101275, doi:10.1016/j.smim.2019.04.003 (2018).

56 Bo, T. H., Dysvik, B. & Jonassen, I. LSimpute: accurate estimation of missing values in microarray data with least squares methods. Nucleic Acids Res 32, e34, doi:10.1093/nar/gnh026 (2004).

57 Liao, Y., Smyth, G. K. & Shi, W. The R package Rsubread is easier, faster, cheaper and better for alignment and quantification of RNA sequencing reads. Nucleic Acids Res 47, e47, doi:10.1093/nar/gkz114 (2019).

58 Love, M. I., Huber, W. & Anders, S. Moderated estimation of fold change and dispersion for RNA-seq data with DESeq2. Genome Biol 15, 550, doi:10.1186/s13059-014-0550-8 (2014).

59 Wu, D. et al. ROAST: rotation gene set tests for complex microarray experiments. Bioinformatics 26, 2176–2182, doi:10.1093/bioinformatics/btq401 (2010).

60 Perales-Paton, J. et al. vulcanSpot: a tool to prioritize therapeutic vulnerabilities in cancer. Bioinformatics 35, 4846–4848, doi:10.1093/bioinformatics/btz465 (2019).

61 Bankhead, P. et al. QuPath: Open source software for digital pathology image analysis. Scientific reports 7, 16878, doi:10.1038/s41598-017-17204-5 (2017).

62 Dimri, G. P. et al. A biomarker that identifies senescent human cells in culture and in aging skin in vivo. Proc Natl Acad Sci U S A 92, 9363–9367 (1995).

